# A revised view on the evolution of glutamine synthetase isoenzymes in plants

**DOI:** 10.1101/2021.11.08.467771

**Authors:** José Miguel Valderrama-Martín, Francisco Ortigosa, Concepción Ávila, Francisco M. Cánovas, Bertrand Hirel, Francisco R. Cantón, Rafael A. Cañas

**Affiliations:** Grupo de Biología Molecular y Biotecnología, Departamento de Biología Molecular y Bioquímica, Universidad de Málaga, Campus Universitario de Teatinos, 29071-Málaga, Spain; Institut National de Recherche pour l’Agriculture, l’Alimentation et l’Environnement (INRAE), Centre de Versailles-Grignon, RD 10, 78026 Versailles cedex, France; Integrative Molecular Biology Lab, Universidad de Málaga, Campus Universitario de Teatinos, 29071-Málaga, Spain

**Keywords:** Glutamine synthetase, phylogeny, plant evolution, nitrogen metabolism, new gene classification

## Abstract

Glutamine synthetase (GS) is a key enzyme responsible for the incorporation of inorganic nitrogen in the form of ammonium into the amino acid glutamine. The genes encoding GS are among the oldest existing genes in living organisms. In plants, two groups of functional GS enzymes are found: eubacterial *GSIIb* (*GLN2*) and eukaryotic *GSIIe* (*GLN1*/*GS*). Phylogenetic analyses have shown that the *GLN2* group originated from bacteria following horizontal gene transfer. Only *GLN1/GS* genes are found in vascular plants, which suggests that they are involved in the final adaptation of plants to terrestrial life. The present phylogenetic study reclassifies the different GS of seed plants into three clusters: *GS1a, GS1b* and *GS2*. The presence of genes encoding GS2 has been expanded to Cycadopsida gymnosperms, which suggests the origin of this gene in a common ancestor of Cycadopsida, Ginkgoopsida and angiosperms. *GS1a* genes have been identified in all gymnosperms, basal angiosperms and some Magnoliidae species. Previous studies in conifers and the gene expression profiles obtained in ginkgo and magnolia in the present work could explain the absence of GS1a in more recent angiosperm species (e.g., monocots and eudicots) due to the redundant roles of GS1a and GS2 in photosynthetic cells. Altogether, the results provide a better understanding of the evolution of plant GS isoenzymes and their physiological roles, which is valuable for improving crop nitrogen use efficiency and productivity.

## INTRODUCTION

Glutamine synthetase (GS, EC 6.3.1.2) catalyzes the incorporation of ammonium into glutamate using ATP to produce glutamine while releasing Pi and ADP (Heldt and Piechulla, 2011). GS is an enzyme of major importance, as it represents the main, if not the only, mechanism incorporating inorganic nitrogen (N) into organic molecules in virtually all living organisms (Shatters et al. 1989). It has been suggested that the genes encoding GS are not only one of the oldest genes in the evolutionary history (Kumada et al. 1993) but also represent an excellent “molecular clock”, which can be used to perform phylogenetic studies (Pesole et al. 1991).

Three GS superfamilies have been identified, namely, GSI, GSII and GSIII, with the corresponding proteins characterized by different molecular masses, different numbers of subunits and their occurrence in the three different domains of life (*Bacteria, Archaea* and *Eukarya*) (Ghoshroy et al. 2010). The GSI superfamily was first found in prokaryotes, although its presence in mammals and plants has also been reported (Mathis et al. 2000; Nogueira et al. 2005; Kumar et al. 2017). The GSII superfamily was described as a group characteristic of *Eukarya* and some *Bacteria* such as *Proteobacteria* and *Actinobacteria* (James et al. 2018). However, the nucleotide sequences deposited in public databases indicate that this GS superfamily is also present in *Euryarchaeota*, a phylum of the *Archaea* domain. Finally, the GSIII superfamily is characteristic of bacteria, including cyanobacteria (James et al. 2018), and some eukaryotes, such as diatoms and other heterokonts, suggesting that GSIII is present in the nucleus of early eukaryotes (Robertson et al. 2006). In a number of studies, the hypothesis that these three gene superfamilies appeared prior to the divergence of eukaryotes and prokaryotes has been proposed (Robertson et al. 2006).

In plants, glutamine synthesis is catalyzed by enzymatic proteins belonging to the GSII superfamily. Two main groups of GSII have been shown to occur in the Viridiplantae group, one of eukaryotic origin, GSII (GSIIe), and the other of eubacterial origin, GSII (GSIIb). *GSIIb* genes are the result of horizontal gene transfer (HGT) following the divergence of prokaryotes and eukaryotes, which in turn represent a sister group of γ-proteobacteria *GSII* (Tateno et al. 1994, Ghoshroy et al. 2010). Since N is one of the main limiting nutrients for plant growth and development, the functions and characteristics of GS have been extensively studied in a large number of vascular plant species, and particularly in crops (Plett et al. 2017; Mondal et al. 2021). It is generally indicated that angiosperms contain two groups of nuclear genes encoding GSIIe represented by cytosolic GS (GS1) and plastidic GS (GS2), each playing distinct physiological roles (Ghoshroy et al. 2010; Hirel and Krapp 2021). GS2 is generally encoded by a single gene, whereas GS1 is encoded by a small multigene family (Cánovas et al. 2007; James et al. 2018). Phylogenetic analyses suggest that *GS2* probably evolved from *GS1* gene duplication (Biesiadka and Legocki 1997) that diverged from a common ancestor 300 million years ago (mya). Therefore, this gene duplication probably occurred before the divergence of monocotyledons and dicotyledons (Bernard and Habash 2009). Interestingly, the gene encoding GS2 is present in the gymnosperm *Ginkgo biloba* (García-Gutiérrez et al. 1998; Guan et al. 2016). This gene is absent in all the other gymnosperms examined thus far, including conifers (Coniferopsida) and gnetales (Gnetopsida), in which the gene encoding GS2 has not been found in their genomes (Birol et al. 2013; Nystedt et al. 2013; Neale et al. 2014; Zimin et al. 2014; Stevens et al. 2016; Neale et al. 2017; Wan et al. 2018; Kuzmin et al. 2019; Mosca et al. 2019; Scott et al. 2020). Furthermore, it seems to also be absent in cycas (Cycadopsida), since the GS2 protein was not detected in western blot analyses (Miyazawa et al. 2018).

In both angiosperm and gymnosperm plants, the synthesis and relative activity of the different GS isoforms are regulated in a species-specific manner but also according to a plant’s developmental stage, tissue, N nutritional status, and to the environmental conditions (Cánovas et al. 2007; Bernard and Habash 2009, Mondale et al. 2021). Consequently, each GS isoform plays a different role during N assimilation and N remobilization throughout a plant’s life cycle (Thomsen et al. 2014; Hirel and Krapp 2021). GS2 predominates in photosynthetic tissues such as leaf mesophyll cells in order to assimilate the ammonium generated from nitrate reduction and released during photorespiration (Wallsgrove et al. 1987; Blackwell et al. 1987; Tegeder and Masclaux-Daubresse 2017). In contrast, GS1 is present in almost all plant organs and tissues (Lea and Miflin 2018). Cytosolic GS isoforms are mostly involved in primary N assimilation in roots and N remobilization and translocation in shoots (Thomsen et al. 2014). As such, it has been shown that they play a key role during plant growth and development, notably for biomass and storage organ production (Xu et al. 2012; Krapp 2015; Havé et al. 2017; Amiour et al. 2021). In conifers, due to the absence of GS2, studies have focused on GS1a and GS1b, which are each encoded by a single gene. These two cytosolic isoforms of GS also exhibit distinct molecular and kinetic properties (Ávila-Sáez et al. 2000; de la Torre et al. 2002). GS1a has been proposed to fulfill the same function as GS2 in angiosperms due to its close relationship with chloroplast development and to the presence of ammonium arising from photorespiration. This hypothesis was also supported by the fact that the gene encoding GS1a is expressed in photosynthetic organs, notably in chlorophyllous parenchyma cells (Ávila et al. 2001), and that its expression is also upregulated in the presence of light (Cantón et al. 1999; Gómez-Maldonado et al. 2004). Moreover, GS1b is phylogenetically and functionally more related to the cytosolic isoforms of GS in angiosperms than to those of GS1a in conifers (Ávila-Sáez et al. 2000; Cánovas et al. 2007).

In this work, the increasing number of plant genome sequences made available in public databases were gathered to perform a deep phylogenetic analysis of the GSII family. The present study includes representative GS sequences from the entire plant evolutionary spectra, including those from monocot and dicot angiosperms and a number of model species that were representative of other taxa. This new phylogenetic study allowed us to propose a revised classification and nomenclature for the different GS isoforms in seed plants. In addition, GS gene expression experiments were conducted in *G. biloba, Magnolia grandiflora* and *Pinus pinaster* in order to strengthen the results obtained in the GS1a phylogeny.

## RESULTS

A total of 168 nucleotide sequences from coding regions (CDS) and the corresponding protein sequences of the genes encoding GSII from 45 different Viridiplantae species were retrieved from different public databases or assembled using next-generation sequencing (NGS) data from the Sequence Read Archive (SRA) database (Table S1). Additionally, *Escherichia coli glnA* (GSI) was used as an external group. The sequences and species analyzed cover the evolutionary history of Viridiplantae and included members of the main Viridiplantae clades unless the sequences were not available in the public databases. These sequences were used to perform phylogenetic analyses to assess GSII evolution in Viridiplantae. The final names of the sequences were assigned depending on phylogenetic analyses (Figures 1 and 2, Table S1). The GSIIb sequences were named GLN2 following the nomenclature of GLN2, the gene encoding GS in *Chlamydomonas reinhardtii*. The GSIIe group, which corresponds to species older than the Embryophyta, was named GLN1. The sequences from Embryophyta species were named GS1, except those included in the group of the Spermatophyta GS2 sequences.

**Figure 1.**
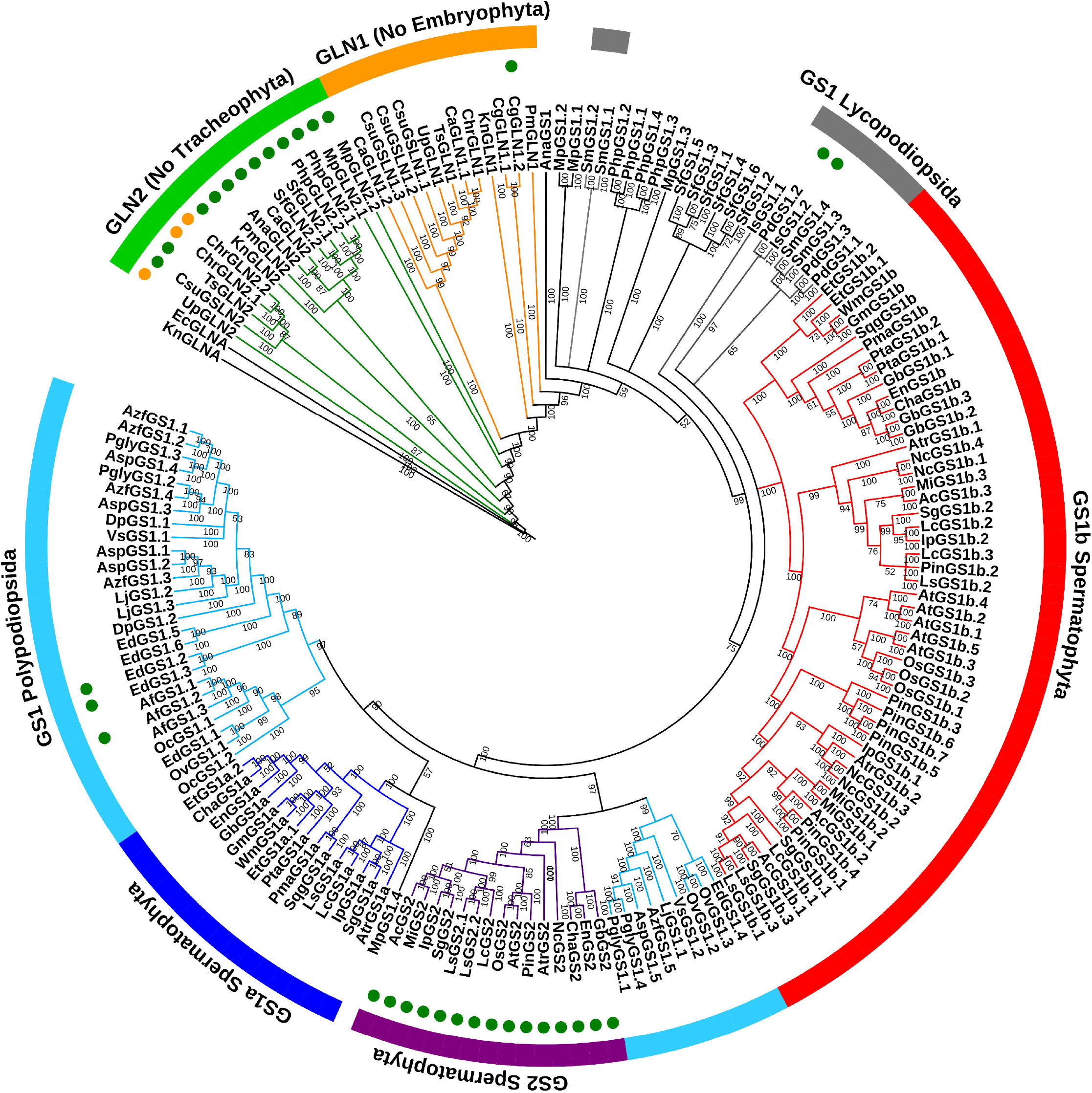
Phylogenetic tree of plant GS nucleotide sequences obtained following a Bayesian analysis. The first two letters of the sequence names correspond to genera and species listed in Table S1. Green circles highlight the sequences exhibiting a predicted plastidic localization. Orange circles highlight the sequences exhibiting a predicted mitochondrial localization. Branch lengths are not presented.

**Figure 2.**
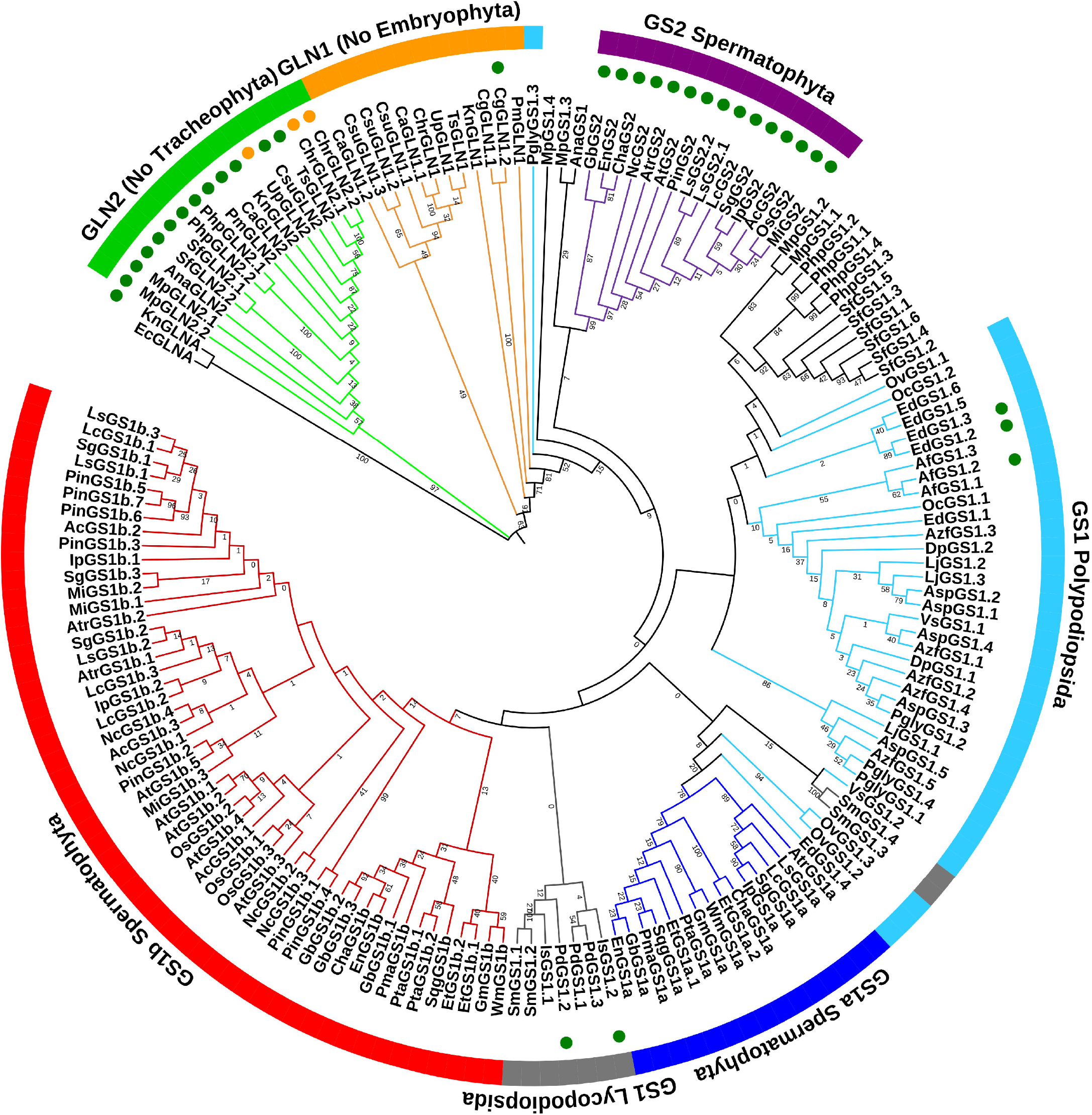
Protein phylogenetic tree of plant GS protein sequences following a Maximum Likelihood analysis. The first two letters of the sequence names correspond to genera and species listed in Table S1. Green circles highlight the sequences with a predicted chloroplastic localization. Orange circles highlight the sequences with a predicted mitochondrial localization. Branch lengths are not presented.

The phylogenetic studies were conducted with nucleotide sequences using Bayesian analyses (Figures 1 and S1). For the protein sequences, a maximum likelihood approach was used (Figures 2 and S2). The results were similar for the main GS groups whether the nucleotide or the protein sequences were analyzed. The *KnGLNA* sequence of *Klebsormidium nitens*, a charophyte green algae, was the most divergent GS close to the outer sequence *EcGLNA* (GSI) of *E. coli* (Figures S1 and S2). Both sequences were very distant from the other plant GS genes (mean length 2.0756), with a node/branch probability of 1 in the Bayesian analysis.

The first cluster contained all GLN2 (GSIIb) sequences. Notably, no GLN2 sequence was identified in vascular plants (Tracheophyta). In the protein sequence analyses, we observed that the GLN2 cluster shared a common origin. In addition, different subgroups for GLN2 were identified in the nucleotide sequence clustering analyses. These GLN2 subgroups were distant from the other plant GS sequences (GSIIe; mean length 0.7142) (Figures S1 and S2). For both the GLN2 nucleotide and protein sequences, the node/branch probability/bootstrap was high (>0.65). In all GLN2 sequences, we found a predicted localization in the chloroplast when using the TargetP software except for UpGLN2, ChrGLN2.1 and ChrGLN2.2 for which we found mitochondrial localization (Figures 1 and 2, Table S1).

The most ancient GSIIe plant sequences were named GLN1, and in both analyses, they were distributed in a main group containing four nonclustered sequences, including *KnGLN1* (Klebsormidiophyceae class), *CgGLN1.1* and *CgGLN1.2* (Coleochaetophyceae class) and *PmGLN1* (Zygnemophyceae class). These four sequences were in an intermediary position between the main GLN1 cluster and the other GS sequences from land plants. A predicted chloroplast localization was only found for CgGLN1.2 (Figures 1 and 2).

Three clusters were identified in seed plants (Spermatophyta), including plastidic GS2 and gymnosperm GS1a-like and GS1b-like sequences (Figures 1 and 2). Interestingly, within the GS2 group, three sequences from gymnosperms were identified (*GbGS2, ChaGS2* and *EnGS2*). They corresponded to a ginkgo and two Cycadopsida sequences. The GS1a-like group contained known GS1a sequences from gymnosperms and GS1 from basal angiosperms and from some Magnoliidae except Ranunculales, Proteales, Liliopsida and Eudycotyledon species. However, an ortholog of the *GS1a* gene was not identified in the genome of *Piper nigrum*, a Magnoliidae species from the Piperales order (Hu et al. 2019). Finally, the GS1b-like cluster contained GS1b found in gymnosperms and the GS1 enzymes previously characterized in angiosperms.

The phylogeny of GS from Anthocerotophyta, Marchantiophyta, Bryophyta, Lycopodiopsida and Polypodiopsida was more complex than that of Spermatophyta, especially when the protein sequences were analyzed (Figures 1 and 2). However, with the cognate gene sequences, Anthocerotophyta, Marchantiophyta, Bryophyta, and Lycopodiopsida were in a basal position compared to the other Embryophyta species. Such a distribution corresponded to the expected evolutionary relationships between plant species, except for the outlier sequence *MpGS1.4* that was found in the GS1a-like cluster (Figure 1). Fern (Polypodiopsida) GS genes were grouped into two clusters. The first one contained most of the nucleotide sequences linked to the GS1a-like sequences, and the second one was grouped with GS2 and was composed of *EdGS1.4, OvGS1.2, OvGS1.3, VsGS1.1, LjGS1.1, AzfGS1.5, AspGS1.5, PglyGS1.1* and *PglyGS1.4*. For these two clusters, the mean probabilities were very high (0.9 and 0.97, respectively) when Bayesian analysis was used (Figure 1, Figure S1).

In contrast, the phylogenetic relationships with the protein sequences were unclear because of a different cluster distribution and the occurrence of outlier sequences such as PglyGS1.3 and MpGS1.4 (Figure 2). Most of the Marchantiophyta and Bryophyta GS enzymes were grouped with most of the Polypodiopsida sequences, although MpGS1.3 and AnaGS1 clustered with the GS2 from Spermatophyta. This group of GS proteins was closer to that of the Spermatophyta GS1 compared to GS2, even though the node/branch bootstraps were very low (<1). The Lycopodiopsida GS enzymes were grouped together with GS1b-like protein sequences even though the node/branch bootstrap was also very low (<1). Nevertheless, two sequences (SmGS1.3 and SmGS1.4) were grouped with the Spermatophyta GS1a cluster together with four Polypodiopsida sequences (EdGS1.4, OvGS1.2, OvGS1.3 and VsGS1.2) (Figure 2).

As expected, for all GS2, the presence of a signal peptide which allows the targeting of the protein to the chloroplast was predicted (Figures 1 and 2, Table S1). Only five of the remaining Embryophyta proteins were predicted to be localized in the chloroplast, including IsGS1.2 and PdGS1.2 from Lycopodiopsida species and EdGS1.2, EdGS1.3 and AfGS1.2 from Polypodiopsida species. Moreover, one could observe that these five GS enzymes did not belong to the GS2 cluster (Figures 1 and 2, Table S1).

### Spermatophyta GS gene expression

To further decipher the role of GS1a in ginkgo and in angiosperms, the level of expression of different GS genes was quantified in *P. pinaster, G. biloba* and *M. grandiflora*. Maritime pine (*P. pinaster*) was included in the study because the role and gene expression pattern of GS1a is well established in this gymnosperm, characterized by the absence of a gene encoding GS2 (Cánovas et al. 2007). The gymnosperm *G. biloba* and the angiosperm *M. grandiflora* were also studied because they possess GS genes corresponding to the three main Spermatophyta GS groups (GS1a, GS1b and GS2).

In maritime pine seedlings, the profiles of *PpGS1a* and *PpGS1b* gene expression were analyzed under different light/dark regimes (Figure 3): germination with a light/dark (L/D) cycle (16 hours of light and 8 hours of darkness), continuous darkness, and two opposite nychthemeral regimes (from light to dark and from dark to light). *PpGS1a* was mainly expressed in the needles irrespective of the light/dark regime and in the stem only during the L/D cycle. In roots, the *PpGS1a* expression level was at the limit of detection under the four different light/dark conditions. In the needles, *PpGS1a* reached the highest level of expression in the dark-light transition, even though it was slightly lower under the L/D cycle. Compared to these two conditions, the *PpGS1a* expression level in the needles was at least five times lower when the plants were placed under continuous darkness and approximately twice lower following a light-dark transition. In contrast, *PpGS1b was* expressed in all three organs. In the needles and in the roots, its expression level was significantly higher only during the light-dark transition. In the stem, the *PpGS1b* expression level was the highest when the seedlings were grown under the L/D cycle.

**Figure 3.**
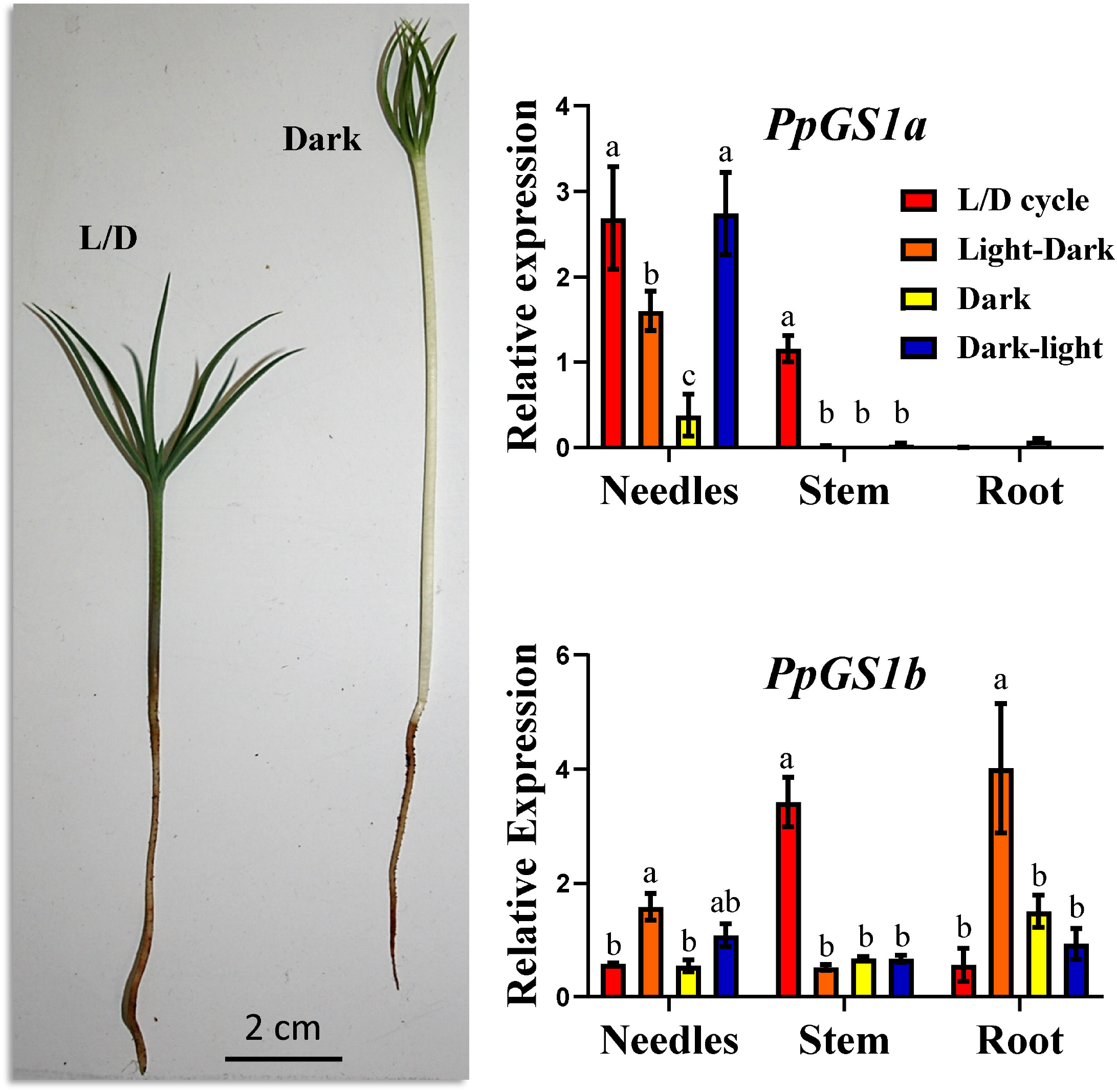
*PpGS1a* and *PpGS1b* gene expression in *Pinus pinaster* seedlings grown under different light regimes. L/D cycle (16h light /8h dark photoperiod, red bars), light photoperiod to continuous darkness transition (orange bars), continuous darkness (yellow bars) and complete darkness to light photoperiod transition (blue bars). Significant differences were determined using Two-way ANOVA that compare the mean for each condition with the mean of the other condition in the same organ. Letters above the columns indicate significant differences when a Tukey’s post-hoc test (*p* < 0.05) was applied.

In *G. biloba* seedlings, *GbGS1a, GbGS2* and *GbGS1b (1 to 3)* expression levels were quantified during the L/D cycle and when plants were placed under continuous darkness (Figure 4A). The absence of leaves in *G. biloba* seedlings germinated under continuous darkness did not allow us to quantify the level of GS gene expression in this organ (García-Gutiérrez et al. 1998). Therefore, light/dark transition experiments were carried out using the leaves of one-year-old *G. biloba* plants (Figure 4B). The amount of *GbGS2 t*ranscripts was very low both in the stems and roots when the seedlings were grown under L/D or continuous darkness conditions. In contrast, under these two conditions, the *GbGS2* expression level was at least 20-fold higher in the leaves. Although the amount of *GbGS1a* transcripts was higher in the leaves than in the other organs, it was four times lower than that of *GbGS2*. The three genes encoding *GbGS1b* were expressed at a higher level in the stems and roots than in the leaves. Two significant correlations were found: between the expression levels of *GbGS1a* and *GbGS2* (0.9) and those of *GbGS1b.1* and *GbGS1b.2* (0.89).

**Figure 4.**
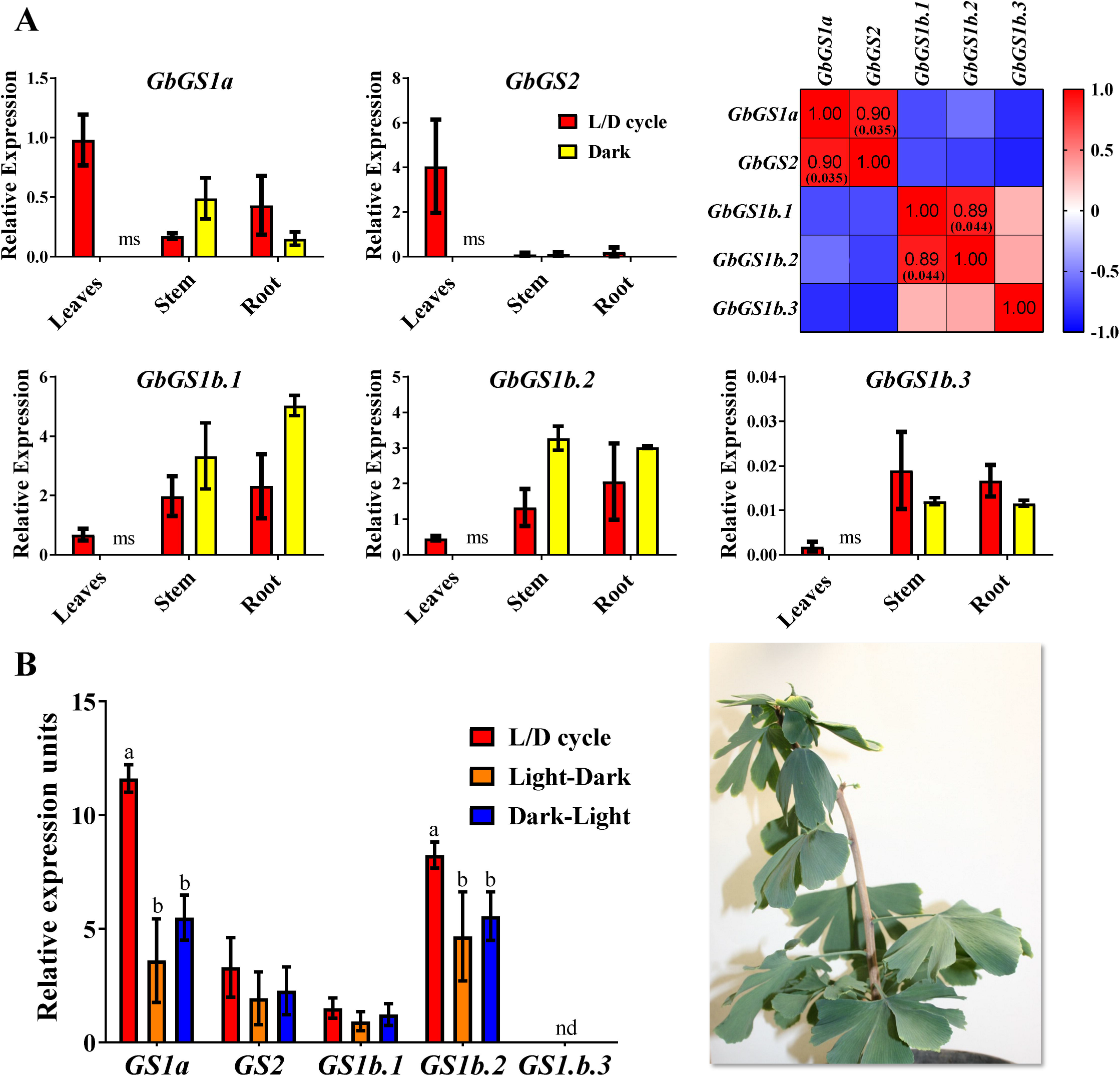
GS gene expression in *Ginkgo biloba* seedlings grown under different light regimes. Panel **A**. One-month old seedlings under L/D cycle: 16h light /8h dark photoperiod (red bars) and continuous darkness (yellow bars). A Pearson correlation test was applied to the expression level of the different GS genes to quantify their relationship indicated in the red squares. The significant *p*-values (< 0.05) for the Pearson coefficient are indicated in brackets. Panel **B**. One-year old seedlings under L/D cycle (16h light /8h dark photoperiod, red bars), light photoperiod to continuous darkness transition (orange bars) and complete darkness to light photoperiod transition (blue bars). Significant differences were determined using a two-way ANOVA to compare the mean for each growth condition with the mean of the other conditions in the same organ. Letters above the columns indicate significant differences based on a Tukey’s post-hoc test (*p* < 0.05). nd= non-detected; ms= missing sample.

When fully expanded leaves were used, *GbGS1a* exhibited the highest level of expression compared to all the other GS genes during the L/D cycle. The pattern of *GbGS2* gene expression was similar to that of *GbGS1a*, although the transcript accumulation was three times lower. Transcripts for *GbGS1b.3* were not detected irrespective of the light/dark regime (Figure 4B).

*M. grandiflora* seedlings were also exposed to different light treatments to study the GS gene expression pattern in this species (Figure 5). The transcripts of *MgGS1a* and *MgGS2* were more abundant in leaves than in stems and roots, and their amounts were similar for *MgGS1a* in the L/D cycle and light-dark treatments. A very low level of expression was obtained for *MgGS1a* when seedlings were placed under continuous darkness. Its level of expression was approximately 4-fold lower than that of the L/D and light/dark treatments following a dark-light transition. *MgGS2* and *MgGS1a* exhibited a similar pattern of transcript accumulation, except that for the former, there was a significant decrease in the light-dark treatment and an increase during the transfer from dark to light. *MgGS1b.1* was the gene exhibiting the highest level of expression compared to all the other genes encoding GS. Its pattern of expression in the different organs was similar to that of *MgGS2*. However, the amounts of *MgGS1b.1* transcripts were much higher in the stems, notably in the L/D conditions, and in the roots. *MgGS1b.2* expression levels were similar in the three organs. No marked differences between the light and dark treatments were observed for this gene. *MgGS1b.3* transcript accumulation was similar irrespective of the organ and light/dark regimes, except in the stem, in which it was much higher during the L/D cycle. As shown in Figure 5, only four significant correlations were found between the expression level of the gene encoding GS in magnolia, where the highest correlation was between *MgGS1a* and *MgGS2* (0.92).

**Figure 5.**
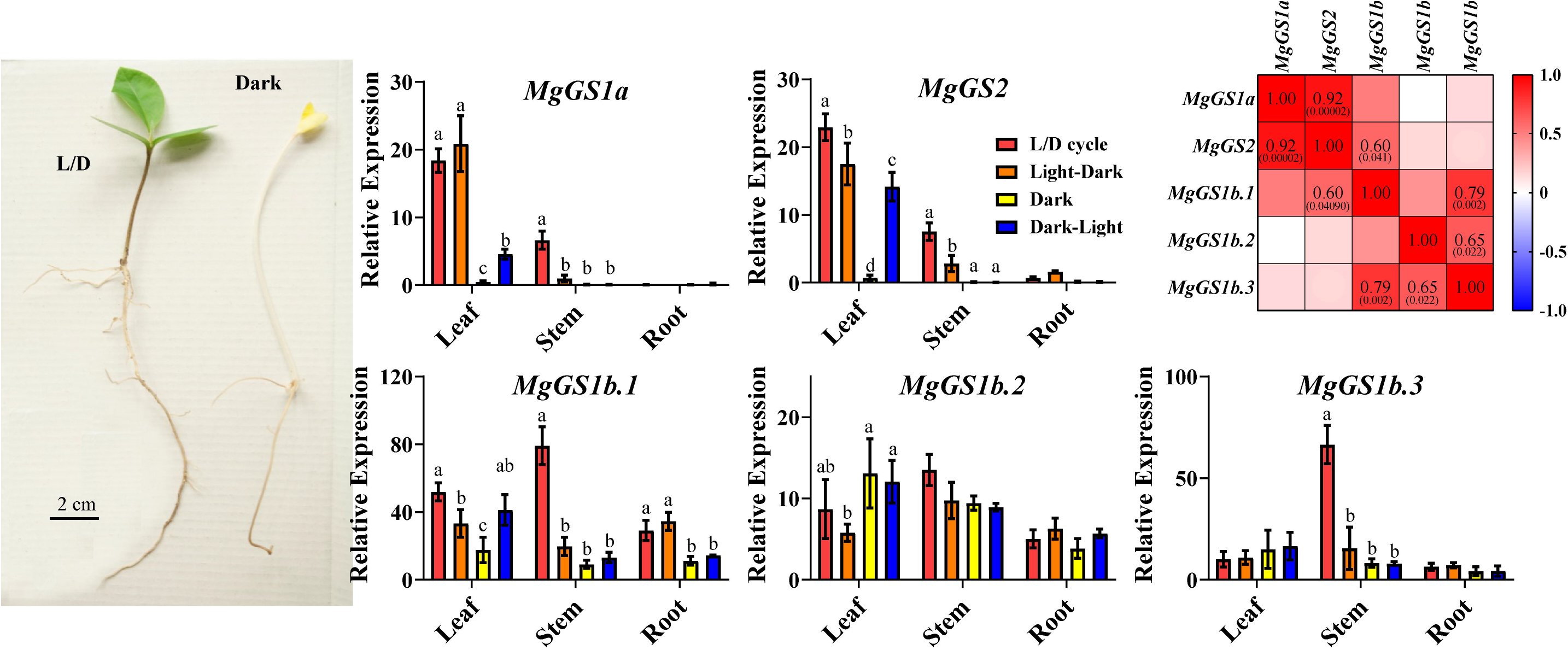
GS gene expression in *Magnolia grandiflora* seedlings grown under different light regimes. L/D cycle (16h light /8h dark photoperiod, red bars), light photoperiod to continuous darkness transition (orange bars), continuous darkness (yellow bars) and complete darkness to light photoperiod transition (blue bars). Significant differences were determined using Two-way ANOVA that compare the mean for each condition with the mean of the other condition for the same organ. Letters above the columns indicate significant differences when a Tukey’s post-hoc test (*p* < 0.05) was applied. Correlations between the expression levels of the different genes expressed in *M. grandiflora* using a Pearson Correlation test. The significant *p*-value (< 0.05) are indicated in brackets.

## DISCUSSION

In all plant species, each GS isoenzyme plays a key role either in primary N assimilation or N recycling, as most of the N-containing molecules required for growth and development are derived from glutamine, the product of the reaction catalyzed by this enzyme. Throughout evolution, such important metabolic functions are subjected to a high selective pressure, which made GS particularly suitable for plant phylogenetic analyses. However, our knowledge of the phylogeny of a number of GS isoenzymes gathered in the evolutionary group called GSII remains limited. The aim of the present study was thus to improve our knowledge on the classification and phylogeny of the Viridiplantae GSII. This study was performed using the corresponding gene sequences belonging to the main clades that are representative of plant evolution and covering a large portfolio of species, not limited to the model and crop angiosperms.

In the present investigation, the resulting phylogenetic analysis agreed with previous studies in which two main groups of plant GSII encoded by nuclear genes were identified, namely, GSIIb (GLN2) and GSIIe (GLN1/GS) (Figures 1 and 2). An HGT event from eubacteria was previously proposed as the more parsimonious process for the emergence of the GLN2 group (Tateno et al. 1994; Ghoshroy et al. 2010). This hypothesis is further supported by the fact that we identified a signal peptide in all the GLN2 that allows the targeting of the proteins to organelles such as plastids and mitochondria (Table S1). Interestingly, genes encoding GLN2 were not identified in vascular plant (tracheophyte) species. Therefore, GLN2 seemed to be lost, coinciding with the final adaptation of plants to land habitats. Such an adaptive mechanism notably included the development of vascular structures for assimilate transport and the presence of lignin involved in plant stature (Raven 2018; Renault et al. 2019). In fact, the massive production of lignin, a metabolic feature of vascular plants, was enabled by a deregulation of phenylalanine biosynthesis that occurred at some point during the evolution of nonvascular plants and tracheophytes (El-Azaz et al. 2021). These developmental and regulatory processes could also be related to the selection of the GLN1/GS genes in the most ancient vascular plants. Thus, it was hypothesized that GLN1/GS isoenzymes are involved 1) in the synthesis of the transport of glutamine and derived amino acids (Bernard and Habash 2009), and 2) in the production of monolignols used as precursors for lignin biosynthesis. As lignin represents one of the main sinks for the photosynthetic carbon assimilated by the plant, high levels of GS activity are thus required to assimilate the large amounts of ammonium released during the reaction catalyzed by the enzyme phenylalanine ammonia-lyase (Pascual et al. 2016). The phylogenetic analyses performed in the present study suggested that the group represented by GLN1 isoenzymes can be considered the starting point for the evolution of the most recent genes encoding GS in plants.

The phylogeny of the ancient Embryophyta clades (Anthocerotophyta, Marchantiophyta and Bryophyta) suggested that the current GS subgroups in Spermatophyta clades were not established in nonvascular land plants (Figures 1 and 2). Curiously, GLN2 was also found in these three clades, which could be the result of a stable situation related to the interaction of genotypic and environmental conditions during the expansion of this group of plants. However, different clustering of GS in Lycopodiopsida and Polypodiopsida was observed between gene and protein phylogenetic trees, resulting in an unclear phylogenetic relationship (Figures 1 and 2). This finding suggests that during plant evolution, there was an active adaptation process as the result of changes in environmental conditions such as the increase in the O_2_/CO_2_ ratio (Renault et al. 2019) and the loss of the *GLN2* gene. According to this hypothesis, several Lycopodiopsida and Polypodiopsida GS protein sequences contain a predicted transit peptide allowing its import into plastids (Table S1). Therefore, under an oxygen-enriched atmosphere, the occurrence of plastidic GS seems to be beneficial for the plant, in turn leading to positive selection.

We also refined the classification of GS in seed plants (Spermatophyta), leading to the identification of three distinct clusters. One was the well-known group of genes from angiosperms encoding plastidic GS (GS2) (see Hirel and Krapp 2021, for a review). The two other clusters contained the genes encoding cytosolic GS (GS1). In one of them, there were all the GS1 isoenzymes classically found in angiosperms and the GS1b from gymnosperms (Cánovas et al. 2007; Bernard and Habash 2009). The third group included the GS1a from gymnosperms, including ginkgo, (Cantón et al. 1993, Ávila-Sáez et al. 2000) and different GS1 from basal angiosperms and some Magnoliidae species. We thus propose to modify the nomenclature of GS from spermatophyte species into three types of genes, namely, GS1a, GS1b and GS2. Consequently, GSIIe can be used as a good phylogenetic marker in seed plants, since the presence or absence of the different GS gene groups is characteristic of the main taxa.

Surprisingly, searches for GS sequences in the NGS data from public databases allowed to identify genes encoding GS2 in Cycadopsida species, contrary to previous findings (Miyazawa et al. 2018). Such a finding was experimentally confirmed by cloning a cDNA encoding GS2 from *Cycas revoluta* (MZ073670). The obtained sequence of the cloned GS2 cDNA from *C. revoluta* validated the assembly of the *Cycas hainanensis* sequence using public NGS data (Figures S3 and S4). Consequently, this result demonstrated that there are more plant clades that possess GS2, which forms a new perspective on the evolution of GS2. In line with such a finding, recent phylogenomic studies showed that Cycadopsida and Ginkgoopsida formed a monophyletic group (Wu et al. 2013; Li et al. 2017; One Thousand Plant Transcriptomes Initiative 2019). The presence of the genes encoding GS2 in both clades and its absence in the other gymnosperm clades (Coniferopsida and Gnetopsida) supports this taxonomic classification.

Based on our phylogenetic analysis, two hypotheses can be proposed concerning GS2 emergence and evolution: 1) A two-event evolutionary process in which the gene encoding GS2 arose from a common ancestor of gymnosperms and angiosperms (Figure 6A). GS1 from Polypodiopsida, which is more closely related to GS2, could have been the origin of the plastid isoform following a specialization process that included the addition of a sequence that allowed the protein to be addressed into the plastids. This hypothesis implies that there was a second genetic event consisting of the loss of GS2 in the common ancestor of Coniferopsida and Gnetopsida plants. 2) A single event during which GS2 sequences emerged from a common ancestor of Cycadopsida/Ginkgoopsida and angiosperm clades, leaving Coniferopsida and Gnetopsida without this gene (Figure 6B). Although gymnosperms are considered a monophyletic clade that is sister to angiosperms (One Thousand Plant Transcriptomes Initiative 2019), the single-event hypothesis about GS2 evolution is more parsimonious and suggests a revision of the phylogenetic relationships of the Cycadopsida/Ginkgoopsida clade with angiosperms.

**Figure 6.**
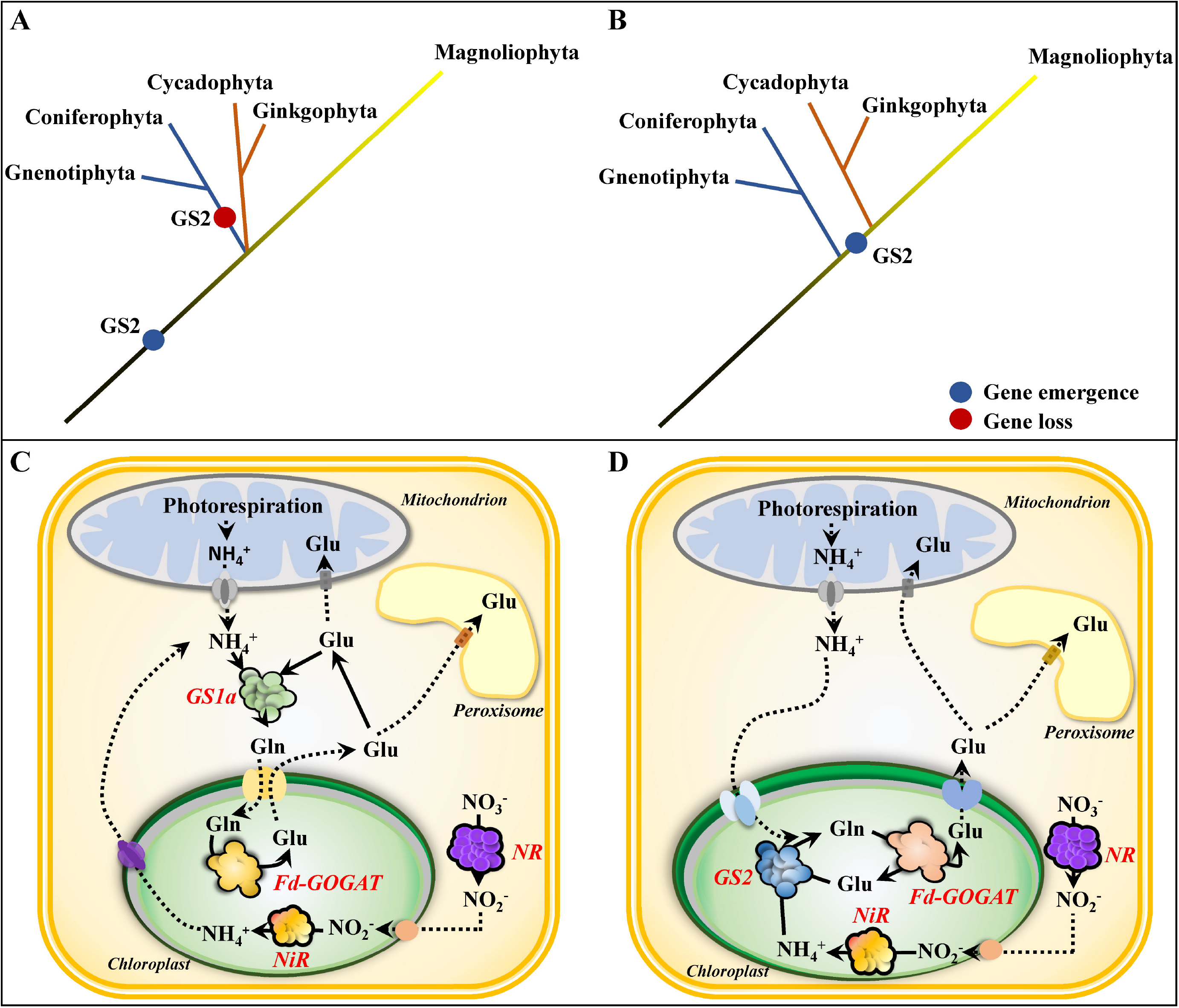
Schematic representation of GS2 emergence hypotheses. **A**. Two-event hypothesis. **B**. Single-event hypothesis. **C**. Simplified metabolic pathways in a photosynthetic cell in which ammonium assimilation is catalyzed by GS1a. **D**. Metabolic pathway of a photosynthetic cell in which ammonium assimilation is catalyzed by GS2. GS, glutamine synthetase. Fd-GOGAT, ferredoxin dependent glutamate synthase. NR, nitrate reductase. NiR, nitrite reductase.

The occurrence of a gene encoding GS1a has never been previously described in ginkgo, basal angiosperms, and a number of Magnoliidae species. The physiological function of GS1a was extensively studied in conifers because it compensates for the lack of GS2. Although GS1a is a cytosolic form of the enzyme, its light-dependent expression level is also associated with chloroplast development, photorespiration and N assimilation and recycling in photosynthetic tissues (Cánovas et al. 2007). One can hypothesize that, even though they are located in different cellular compartments, GS1a and GS2 play redundant roles in photosynthetic cells, which could explain the disappearance of GS1a in the most recent angiosperm species. The light dependence and organ gene expression of GS1a and GS2 in *P. pinaster* (Figure 3), *G. biloba* (Figure 4) and *M. grandiflora* (Figure 5) strengthened the previous hypothesis that GS1a fulfills the function of GS2 (Cantón et al. 1999; Ávila et al. 2001; Gómez-Maldonado et al. 2004). Interestingly, one of the two Cys residues involved in the redox modulation of GS2 activity (e.g., C306 in *Arabidopsis*) was conserved in all the GS1a proteins (Cantón et al., 1993; Choi et al. 1999; Miyazawa et al. 2018). We also observed that this residue was conserved in all the GS1a and GS2 protein sequences analyzed in this study and in those from Polypodiopsida (ancient Embryophyta clades), a number of Lycopodiopsida species and in GLN1 sequences (Figure S5). Remarkably, this Cys is absent in all GS1b, suggesting a specific role of this residue in the function of GS2 and GS1a. In angiosperms, a second Cys residue is present in GS2 (e.g., C371 in *Arabidopsis*) and in a number of GLN1 proteins. Moreover, this second Cys residue is not present in ginkgo or Cycadopsida GS2, which suggests that it was acquired by angiosperms during plant evolution (Figure S5).

When the single-event hypothesis is considered, the emergence of GS2 following the loss of GS1a in recent angiosperms indicates that they probably have redundant physiological functions. The Cycadopsida/Ginkgoopsida group emerged at least 270 mya ago (Wu et al. 2013; Li et al. 2017; One Thousand Plant Transcriptomes Initiative 2019) during the Permo-Carboniferous period, when the atmospheric oxygen level rose from 21 to 35%. Such an elevation in oxygen level led to a drastic increase in the oxygenase activity of the photosynthetic enzyme Rubisco, leading to increased photorespiration (Berling and Berner, 2000). The series of events that occurred during the Permo-Carboniferous period could result in the appearance of a plastidic GS isoenzyme (GS2) *via* a positive selection process, allowing for a more efficient reassimilation of ammonium that is released during photorespiration. Mutation of GS2 in a number of species induced lethality under photorespiratory conditions (Wallsgrove et al. 1987; Blackwell et al. 1987; Pérez-Delgado et al. 2015), which is not the case in *Arabidopsis*, as it that can cope with the toxicity of ammonium released from the photorespiratory pathway (Ferreira et al. 2019; Hachiya et al. 2021). Conifers, which are C3 species possessing only cytosolic GS isoenzymes (Figure 6C), are also able to reassimilate the ammonium released during photorespiration. However, the increase in the amount of atmospheric oxygen during the Permo-Carboniferous would imply an increase in nitrification rates, since oxygen is a substrate for nitrification (Ward, 2008), thus leading to an increase in nitrate availability in the rhizosphere. One can therefore hypothesize that such an increase in nitrate availability induced additional evolutionary pressure toward the selection of plastidic GS (GS2), which, in addition to photorespiratory ammonium reassimilation, is also responsible for the assimilation of ammonium in plastids derived from nitrate reduction (Figure 6D) (Hirel and Krapp 2021). Consistent with this hypothesis, it is known that most conifers prefer or tolerate ammonium as an inorganic N source, which can be readily assimilated by cytosolic GS in the absence of GS2. Concerning the process of GS2 selection, the most likely hypothesis is the duplication of genes encoding cytosolic GS leading to functional specialization due to changes such as those in the gene promoter and the addition of a sequence encoding a signal peptide used to import the protein into the chloroplasts (Biesiadka and Legocki 1997, Ávila-Sáez et al. 2000). Gene expression patterns, specific Cys residue conservation and nucleotide sequence-based GS phylogeny suggest that GS1a could be at the origin of GS2 (Figures 1, 3–5).

Finally, we were able to conclude that the group represented by GS1b evolved in a different way than that grouping GS1a and GS2. In ginkgo and angiosperms, GS1b is generally represented by a small multigene family, with each member playing distinct roles either in N assimilation or N recycling, depending on the organ examined (Thomsen et al. 2014). In contrast, there is usually only one gene of GS1a or GS2, strongly suggesting that GS1b genes play nonredundant roles compared to GS1a and GS2 (Ghoshroy et al., 2010; Hirel and Krapp 2021). Related to their different roles, the comparison between the GS1a and GS1b proteins in pine showed distinctive characteristics, such as a higher thermal stability of GS1b (de la Torre et al. 2002).

### Conclusions

The combined phylogenetic analysis and gene expression study presented in this work allowed us to improve our understanding of GS evolution in plants, notably in the Spermatophyta clade. In agreement with previous studies, two distinct groups of GS, GSIIb (GLN2) and GSIIe (GLN1 and GS), were clearly identified (Tateno et al. 1994; Ghoshroy et al. 2010).

An original finding of our study was that in seed plants, GS is represented by three types of genes (GS1a, GS1b and GS2), which allowed us to redefine the nomenclature of GS1 isoenzymes. In particular, the taxa possessing a gene encoding GS1a that was originally represented by conifers (Cantón et al. 1993, 1999), are now expanded to ginkgo, basal angiosperms, and some Magnoliidae species. In contrast, GS1a is not present in the most recent angiosperm species such as monocots and eudicots. Therefore, GS1a represents a new functional group as this cytosolic GS isoenzyme is able to compensate for the lack of GS2 (Ávila et al. 2001).

Unexpectedly, a gene encoding GS2 was identified in Cycadopsida species. This new finding supports that this clade, together with Ginkgoopsida species, represents a monophyletic group (Wu et al. 2013). Concerning the emergence and evolution of GS2, we provide strong lines of evidence that this gene arose in the most recent common ancestor of Cycadopsida, Ginkgoopsida and angiosperms. However, additional analyses including GS gene sequences from ferns will be required to confirm this hypothesis.

Beyond the numerous studies already performed on crops, the present investigation proposes new possibilities toward a better understanding of the evolution of GS genes and their functions in terrestrial plants. Additional studies of GS evolution could help further deciphering the impact of the different GS isoenzymes on N use efficiency to improve plant growth and productivity under a wide range of environmental conditions.

## MATERIALS AND METHODS

### Glutamine synthetase gene sequences and phylogenetic analyses

Nucleotide sequences of the genes encoding plant glutamine synthetase type II (GSII) were obtained from different public databases or assembled from transcriptomic NGS data from SRA database at the National Center for Biotechnology Information (NCBI) and at the European Nucleotide Archive (EBI) (Table S1). The tool employed for sequence search was BLAST (Altschul et al. 1990) using mainly the *tblastn* mode with the sequence of GS1b.1 from *Pinus taeda* as the query. For the assembly of sequences from NGS data, the raw files were uploaded to the web platform Galaxy (https://usegalaxy.org/), which was used to make the transcriptome assemblies (Afgan et al. 2016). The raw reads were quality trimmed with the Trimmomatic software (Bolger et al. 2014). The transcriptome assemblies were conducted with the Trinity assembler (Grabherr et al. 2011) and the GS sequences were identified using BLAST as described above. Database identifiers, names and species for the different GS sequences are presented in Table S1. All nucleotide sequences used in the present work are shown in Dataset S1. The subcellular localization prediction was determined with TargetP (Almagro Armenteros et al. 2019) and LOCALIZER software (Sperschneider et al. 2017).

For Bayesian phylogenetic analysis a dataset of 169 nucleotide sequences encoding GSII from 45 different Viridiplantae species and *glnA* from *Escherichia coli* were aligned with the Muscle software (Edgar, 2004). Position with gaps were deleted, and MRMODELTEST v2.4 was used to find the best fit model among 24 models used to study molecular evolution (Nylander, 2004). The Akaike Information Criterion suggested to use model GTR+I+G. The Bayesian phylogenetic analysis was performed using MRBAYES v3.2.7 (Huelsenbeck and Ronquist, 2001) with two simultaneous runs of 77 million generations for each run, with one cold and three heated chains for each run in which the temperature parameter was set to 0.1. Trees were sampled once every 10,000 generations. The average standard deviation of split frequencies at the end of each run was < 0.01, and the first 25 % of the trees were discarded as burn-in samples. The consensus tree was visualized with the interactive Tree Of Life (iTOL) web tool (Letunic and Bork 2019).

For maximum likelihood analysis the dataset composed of 169 GS protein sequences obtained from the corresponding nucleotide sequences that were used for the Bayesian analyses. The alignment and phylogenetic analysis were conducted using MEGA7 (Kumar et al. 2016). The sequences were aligned with Muscle (Edgar, 2004). Maximum likelihood analyses were carried out using the complete deletion of gaps, the missing data, and the amino acid substitution model Jones-Taylor-Thornton (JTT) (Jones et al. 1992). The Nearest-Neighbor-Interchange (NNI) was used for tree inference. The initial tree was constructed using the NJ/BioNJ method. The phylogeny test was performed using the Bootstrap method with 1,000 replications.

### Plant material

*G. biloba* seeds were obtained from different botanic gardens: Botanische Gärten der Universität Bonn (Bonn, Germany), Botanischer Garten der Universität Bern (Bern, Switzerland), Plantentuin Universteit Gent (Ghent, Belgium) and Arboretum Wespelaar (Wespelaar, Belgium). Ginkgo seeds were stratified for 3 months in vermiculite at 4°C. *P. pinaster* seeds from Sierra Bermeja (Estepona, Spain) (ES20, Ident. 11/12) were obtained from the *Red de Centros Nacionales de Recursos Genéticos Forestales* of the Spanish *Ministerio para la Transición Ecológica y el Reto Demográfico* with the authorization number ESNC87. Pine seeds were imbibed for 72 hours under continuous aeration with an air pump*. M. grandiflora* seeds were obtained from the Parque de la Alameda garden in Málaga (Spain) and from private suppliers. Magnolia seeds were stratified in vermiculite at 4°C for 4 months. All seeds were growth on vermiculite in order to prevent any nutritional effect of the substrate.

Ginkgo, pine, and magnolia seedlings were germinated and grown at 23°C either with a 16h light/8h dark photoperiod and watered once every 3 days with distillated water or under continuous darkness and watered once a week. For light/dark transition experiments, seedlings grown in complete darkness were transferred to a 16h light/8h night photoperiod for 24 hours and seedlings grown with 16h light/8h night were transferred to complete darkness for 24 hours. Leaves, stems, and roots of seedlings were harvested separately. For ginkgo seedlings grown in complete darkness, primary and secondary leaves were not developed, thus only stems and roots were harvested. To study the impact of a light-dark transition on the expression of the different genes encoding GS in ginkgo leaves, one-year old plants with fully developed leaves were used. These plants were first exposed to a 16h light /8h night photoperiod, then to complete darkness for 24 hours and then back to a 16h light /8h night cycle. The different plant samples were immediately frozen in liquid N and stored at −80°C.

For the cloning of *Cycas revoluta GS2*, leaves from a one-year-old plants grown under 16h light/8h night photoperiod were harvested. Leaf tissues were frozen immediately in liquid N and stored at −80°C until further use for RNA extraction.

### RNA extraction and RT-qPCR

Ginkgo and magnolia RNAs were extracted using the Plant/Fungi Total RNA Purification Kit (Norgen Biotek Corp., Thorold, ON, Canada) according to the manufacturer’s instruction manual. Pine and Cycas RNAs were extracted as described by Canales et al. (2012). For the cDNA synthesis, 500 ng of total RNA were used and retrotranscribed using the iScrpt™ Reverse Transcription Supermix (Bio-Rad, Hercules, CA, USA). qPCR was carried out using 10 ng of cDNA and the SsoFast™ EvaGreen® Supermix (Bio-Rad, Hercules, CA, USA). The reaction was carried out in a thermal cycler CFX384™ Touch Real-Time PCR (Bio-Rad, Hercules, CA, USA). The results for maritime pine were normalized using a *saposin-like aspartyl protease* (unigene1135) as a reference gene (Granados et al. 2016). A number of references genes used to study gene expression in maritime pine (Granados et al. 2016) were tested in gingko and magnolia. In this species, the orthologs of maritime pine *saposin-like aspartyl protease* (unigene1135), *myosin heavy chain-related* (unigene13291) and of an RNA binding protein (unigene27526) were selected and used to normalize the expression of the genes encoding GS. In magnolia, the ortholog of an *RNA binding protein* (unigene27526) from maritime pine and *Actin-7* from magnolia (Lovisetto et al. 2015) were selected and used as reference genes. The different primers used for the RT-qPCR experiments are presented in Table S2. Magnolia and ginkgo sequences used to design the primers are listed in Table S3.

A GS2 cDNA from *C. revoluta* was cloned using a PCR product. An iProof™ HF Master Mix (Bio-Rad, Hercules, CA, USA) was used to perform the PCR reaction. Primer sequences were obtained from *C. hainanensis* and presented in Table S2. After the initial denaturation step at 98°C during 1 min, the PCR was conducted during 35 cycles with the following conditions: 10 s at 98°C; 20 s at 60°C and 1 min at 72°C, with a final extension step at 72°C for 5 min. The resulting PCR product was cloned into the pJET1.2 cloning vector (Thermo, Waltham, MA, USA). The sequence of GS2 from *C. revoluta* was submitted to Genbank (MZ073670).

### Statistics

Statistical analyses were performed using Prism 8 (Graphpad, CA, USA). Data obtained from gene expression quantification were analyzed using a multiple comparison Two-Way Anova test. Differences between organs were not analyzed statistically. The Tukey’s test was used as a post hoc test for statistical analysis of the gene expression data. For *G. biloba* and *M. grandiflora* seedlings a Pearson correlation test was also used to evaluate the relationship existing between the expressions of the different *GS* genes. Differences and correlations were considered to be significant when the *p*-value was < 0.05.

## ACKNOWLEDGEMENTS

The authors are very grateful to Dr. Juan Antonio Pérez Claros for his helpful comments on the manuscript and Anett Krämer (Botanische Gärten der Universität Bonn, Germany), Katja Rembold (Botanischer Garten der Universität Bern, Switzerland), Chantal Dugardin (Plantentuin Universteit Gent, Belgium) and Joke Ossaer (Arboretum Wespelaar, Belgium) for the kind supply of ginkgo seeds. We are also very grateful to José Manuel Sánchez Calle and María Dolores Trujillo Gutiérrez for the kind supply of magnolia seeds. This work was supported by Spanish *Ministerio de Ciencia e Innovación*, grant numbers BIO2015-73512-JIN MINECO/AEI/FEDER, UE; RTI2018-094041-B-I00 and EQC2018-004346-P. This work was also supported by *Junta de Andalucía*, grant number P20_00036 PAIDI 2020/FEDER, UE. JMVM was supported by a grant from the Spanish *Ministerio de Educación y Formación Profesional* (FPU17/03517). FO was supported by grants from the *Universidad de Málaga (Programa Operativo de Empleo Juvenil vía SNJG, UMAJI11, FEDER, FSE, Junta de Andalucía*) and BIO-114, *Junta de Andalucía*.

## AUTHORS CONTRIBUTION

JMVM and FO have performed the experiments; FRC and RAC have performed the phylogenetic analysis; JMVM, BH, FRC and RAC have written the manuscript; JMVM, FO and RAC have designed the figures; FMC and CA have made additional contributions and edited the manuscript. RAC, CA, and FMC were responsible of the funding acquisition; FRC and RAC have planned and designed the research.

## DATA AVAILABILITY

The data that support the findings of this study are available from different databases, supporting information and from the corresponding author upon reasonable request.

## SUPPLEMENTAL DATA

**Table S1.** Sequences names, accession numbers, species taxonomy data and putative subcellular localization of the different encoded GS.

**Table S2.** List of primers used for RT-qPCR experiments *CrGS2* cloning.

**Table S3.** Magnolia and ginkgo GS gene sequences used to design the primers used for RT-qPCR experiments.

**Dataset S1.** GS nucleotide sequences used to perform the phylogenetic analyses.

**Dataset S2.** GS protein sequences to perform the phylogenetic analyses.

**Dataset S3.** Detailed results of the Bayesian analysis using the GS gene nucleotide sequences.

**Dataset S4.** Detailed results of the maximum likelihood analysis using the GS protein sequences.

**Figure S1.** Phylogenetic tree obtained following a Bayesian analysis of the GS nucleotide sequences in which the branch lengths are maintained. Green circles correspond a predicted plastid localization of the corresponding protein. Orange circles correspond to a predicted mitochondrial localization.

**Figure S2.** Phylogenetic tree obtained following a Bayesian analysis of the GS protein sequences in which the branch lengths are maintained. Green circles correspond predicted plastid localization. Orange circles correspond to a predicted mitochondrial localization.

**Figure S3.** Multiple sequence alignment of the GS2 protein sequences from *Cycas revoluta* (CrGS2) and *Cycas hainanensis* (ChaGS2).

**Figure S4.** Multiple sequence alignment of the complete coding sequences (CDS) of the genes encoding GS2 from *Cycas revoluta* (*CrGS2*) and *Cycas hainanensis* (*ChaGS2*).

**Figure S5.** Multiple sequence alignment of the protein regions around the Cys residues involved in the redox modulation of GS2 activity, C306 and C371 positions in *Arabidopsis* GS2.

## REFERENCES

Afgan E, Baker D, Batut B, van den Beek M, Bouvier D, Cech M, Chilton J, Clements D, Coraor N, Grüning B, et al. 2018. The Galaxy platform for accessible, reproducible and collaborative biomedical analyses: 2018 update. Nucleic Acids. Res. 46:W537–W544.

Almagro Armenteros JJ, Salvatore M, Emanuelsson O, Winther O, von Heijne G, Elofsson A, Nielsen H. 2019. Detecting sequence signals in targeting peptides using deep learning. Life Sci. Alliance 2:e201900429.

Amiour N, Décousset L, Rouster J, Quenard N, Buet C, Dubreuil P, Quilleré I, Brulé L, Cukier C, Dinant S, et al. 2021. Impacts of environmental conditions, and allelic variation of cytosolic glutamine synthetase on maize hybrid kernel production. Commun Biol. 4:1095.

Altschul SF, Gish W, Miller W, Myers EW, Lipman DJ. 1990. Basic local alignment search tool. J. Mol. Biol. 215:403–410.

Ávila-Sáez C, Muñoz-Chapuli R, Plomion C, Frigerio J, and Cánovas FM. 2000. Two genes encoding distinct cytosolic glutamine synthetases are closely linked in the pine genome. FEBS Lett. 477:237–43.

Ávila C, Suárez MF, Gómez-Maldonado J, and Cánovas FM. 2001. Spatial and temporal expression of two cytosolic glutamine synthetase genes in Scots pine: functional implications on nitrogen metabolism during early stages of conifer development. Plant J. 25:93–102.

Berling DJ, Bermer RA. 2000. Impact of a Permo-Carboniferous high O2 event on the terrestrial carbon cycle. Proc. Nat. Acad. Sci. U.S.A. 97:12428–12432.

Bernard SM, Habash DZ. 2009. The importance of cytosolic glutamine synthetase in nitrogen assimilation and recycling. New Phytol. 182:608–620.

Biesiadka J, Legocki AB. 1997. Evolution of the glutamine synthetase gene in plants. Plant Sci. 128: 51–58.

Birol I, Raymond A, Jackman S.D, Pleasance S, Coope R, Taylor G.A, Yuen MMS, Keeling CI, Brand D, Vandervalk BP, et al. 2013. Assembling the 20 Gb white spruce (*Picea glauca*) genome from whole-genome shotgun sequencing data. Bioinformatics 29 12:1492–1497.

Bolger AM, Lohse M, Usadel B. 2014. Trimmomatic: A flexible trimmer for Illumina Sequence Data. Bioinformatics 30: 2114–2120.

Blackwell RD, Murray AJS, Lea PJ. 1987. Inhibition of photosynthesis in barley with decreased levels of chloroplastic glutamine synthetase activity. J. Exp. Bot. 196:1799–1809.

Canales J, Rueda-López M, Craven-Bartle B, Avila C, Cánovas FM. 2012. Novel insights into regulation of asparagine synthetase in conifers. Front. Plant Sci. 24:3–100.

Cánovas FM, Avila C, Cantón FR, Cañas RA, de la Torre F. 2007. Ammonium assimilation and amino acid metabolism in conifers. J. Exp. Bot. 58:2307–2318.

Cantón FR, García-Gutiérrez A, Gallardo F, de Vicente A, Cánovas FM. 1993. Molecular characterization of a cDNA clone encoding glutamine synthetase from a gymnosperm, *Pinus sylvestris*. Plant Mol. Biol. 22:819–828.

Cantón FR, Suárez MF. Josè-Estanyol M, Cánovas FM. 1999. Expression analysis of a cytosolic glutamine synthetase gene in cotyledons of Scots pine seedlings: developmental, light regulation and spatial distribution of specific transcripts. Plant. Mol. Bio. 404:623–34.

Choi YA, Kim SG, Kwon YM. 1999. The plastidic glutamine synthetase activity is directly modulated by means of redox change at two unique cysteine residues. Plant Sci. 149:175–182.

de la Torre F, García-Gutiérrez A, Crespillo R, Cantón FR, Avila C, Cánovas FM. 2002. Functional expression of two pine glutamine synthetase genes in bacteria reveals that they encode cytosolic holoenzymes with different molecular and catalytic properties. Plant Cell Physiol. 437:802–809.

Edgar R. C. 2004. MUSCLE: multiple sequence alignment with high accuracy and high throughput. Nucleic Acids Res. 32:1792–1797.

El-Azaz J, Cánovas FM, Barcelona B, Avila C, de la Torre F. 2021. Deregulation of phenylalanine biosynthesis evolved with the emergence of vascular plants. Plant Physiol. (forthcoming) doi:10.1093/plphys/kiab454.

Ferreira S, Moreira E, Amorim I, Santos C, Melo P. 2019. *Arabidopsis thaliana* mutants devoid of chloroplast glutamine synthetase (GS2) have non-lethal phenotype under photorespiratory conditions. Plant Physiol. Biochem. 144:365–374.

García-Gutiérrez A, Dubois F, Cantón FR, Gallardo F, Sangwan RS, and Cánovas F. 1998. Two different models of early development and nitrogen assimilation in gymnosperm seedlings. Plant J. 132:187–199.

Granados JM, Ávila C, Cánovas FM, Cañas RA. 2016. Selection and testing of reference genes for accurate RT-qPCR in adult needles and seedlings of maritime pine. Tree Genet. Genomes 12:60–75.

Ghoshroy S, Binder M, Tartar A, Robertson DL. 2010. Molecular evolution of glutamine synthetase II: phylogenetic evidence of a non-endosymbiotic gene transfer event early in plant evolution. BMC Evol. Biol. 10:198.

Gómez-Maldonado J, Avila C, de la Torre F, Cañas R, Cánovas FM, Campbell MM. 2004. Functional interactions between a glutamine synthetase promoter and MYB proteins. Plant J. 39:513–26.

Grabherr MG, Haas BJ, Yassour M, Levin JZ, Thompson DA, Amit I, Adiconis X, Fan L, Raychowdhury R, Zeng Q, et al. 2011. Full-length transcriptome assembly from RNA-seq data without a reference genome. Nat Biotechnol. 29:644–652.

Guan R, Zhao Y, Zhang H, Fan G, Liu X, Zhou W, Shi C, Wang J, Liu W, Liang X, et al. 2016. Draft genome of the living fossil *Ginkgo biloba*. Gigascience 5:49.

Hachiya T, Inaba J, Wakazaki M, Sato M, Toyooka K, Miyagi A, Kawai-Yamada M, Sugiura D, Nakagawa T, Kiba T, et al. 2021. Excessive ammonium assimilation by plastidic glutamine synthetase causes ammonium toxicity in *Arabidopsis thaliana*. Nat. Commun. 12:4944.

Heldt H, Piechulla B. 2011. Plant Biochemestry: fourth edition. San Diego: Academic Press, Elsevier.

Havé M, Marmagne A, Chardon F, Masclaux-Daubresse C. 2017. Nitrogen remobilization during leaf senescence: lessons from *Arabidopsis* to crops. J. Exp. Bot. 68:2513–2529.

Hirel B and Krapp A. 2021. Nitrogen Utilization in Plants I Biological and Agronomic Importance. Encyclopedia of Biological Chemistry III (Third Edition). Editor: Joseph Jez, 1: 127–140. Elsevier. ISBN 9780128220405, doi:10.1016/b978-0-12-809633-8.21265-x.

Hu L, Xu Z, Wang M, Fan R, Yuan D, Wu B, Wu H, Qin X, Yan L, Tan L, et al. 2019. The chromosome-scale reference genome of black pepper provides insight into piperine biosynthesis. Nat. Commun. 10:4702.

Huelsenbeck JP, Ronquist F. 2001. MRBAYES: Bayesian inference of phylogeny. Bioinformatics 17:754–755.

James D, Borphukan B, Fartyal D, Achary VMM, Reddy MK. 2018. Transgenic manipulation of glutamine synthetase: A target with untapped potential in various aspects of crop improvement. In: Gosal SS, Wani SH, editors. Biotechnology of Crop Improvement. Springer International Publishing AG. p. 367–416.

Jones DT, Taylor WR, Thornton JM. 1992. The rapid generation of mutation data matrices from protein sequences. Comput. Appl. Biosci. 8:275–282.

Krapp A. 2015. Plant nitrogen assimilation and its regulation: a complex puzzle with missing pieces. Curr. Opin. Plant. Biol. 25:115–122.

Kumada Y, Benson DR, Hillemann D, Hosted TJ, Rochefort DA, Thompson CJ, Wohlleben W, Tateno, Y. 1993. Evolution of the glutamine synthetase gene, one of the oldest existing and functioning genes. Proc. Nat. Acad. Sci. U.S.A. 90:3009–3013.

Kumar S, Stecher G, Tamura K. 2016. MEGA7: Molecular Evolutionary Genetics Analysis Version 7.0 for Bigger Datasets. Mol. Biol. Evol. 33:1870–1874.

Kumar V, Yadav S, Soumya N, Kumar R, Babu NK, Singh S. 2017. Biochemical and inhibition studies of glutamine synthetase from *Leishmania donovani*. Microb. Pathog. 107:164–174.

Kuzmin DA, Ferachuk SI, Sharov VV, Cybin AN, Makolov SV, Putintseva A, Oreshkova NV, Krutovsky KV. 2019. Stepwise large genome assembly approach: a case of Siberian larch (*Larix sibirica* Ledeb). BMC Bioinformatics 20:37.

Lea PJ, Miflin BJ. 2018. Nitrogen assimilation and its relevance to crop improvement. In: Foyer CH, Zhang H, editors. Annual Plant Reviews. Volume 42. Nitrogen Metabolism In Plants in the Post-Genomic Era. Chichester: Wiley-Blackwell, pp. 1–40.

Letunic I, and Bork P. 2019. Interactive Tree Of Life (iTOL) v4: recent updates and new developments. Nucleic Acids Res. 47: W256–W259.

Li Z, De La Torre AR, Sterck L, Cánovas FM, Avila C, Merino I, Cabezas JA, Cervera MT, Ingvarsson PK, Van de Peer Y. 2017. Single-Copy Genes as Molecular Markers for Phylogenomic Studies in Seed Plants. Genome Biol. Evol. 9:1130–1147.

Lovisetto A, Masiero S, Rahim MA, Mendes MA, Casadoro G. 2015. Fleshy seeds form in the basal Angiosperm *Magnolia grandiflora* and several MADS-box genes are expressed as fleshy seed tissues develop. Evol Dev. 1:82–91.

Mathis R, Gamas P, Meyer Y, Cullimore JV. 2000. The presence of GSI-like genes in higher plants: support for the paralogous evolution of GSI and GSII. J. Mol. Evol. 50:116–122.

Miyazawa SI, Nishiguchi M, Futamura N, Yukawa T, Miyao M, Maruyama TE, Kawahara T. 2018. Low assimilation efficiency of photorespiratory ammonia in conifer leaves. J Plant Res. 131:789–802.

Mondal R, Kumar A, Chattopadhyay SK. 2021. Structural property, molecular regulation and functional diversity of Glutamine Synthetase in higher plants: a data-mining bioinformatics approach. Plant J. (in press). doi: 10.1111/TPJ.15536

Mosca E, Cruz F, Gómez-Garrido J, Bianco L, Rellstab C, Brodbeck S, Csilléry K, Fady B, Flandug M, Fussi B, et al. 2019. A Reference Genome Sequence for the European Silver Fir (*Abies alba* mill.): A Community-Generated Genomic Resource. G3 (Bethesda) 9:2039–2049.

Neale DB, Wegrzyn JL, Stevens KA, Zimin AV, Puiu D, Crepeau MW, Cardeno C, Koriabine M, Holtz-Morris AE, Liechty JD, et al. 2014. Decoding the massive genome of loblolly pine using haploid DNA and novel assembly strategies. Genome Biol. 15:R59.

Neale DB, McGuire PE, Wheeler NC, Stevens KA, Crepeau MW, Careno C, Zimin AV, Puiu D, Pertea GM., Sezen UU, et al. 2017. The Douglas-Fir Genome Sequence Reveals Specialization of the Photosynthetic Apparatus in Pinaceae. G3 (Bethesda) 7:3157–3167.

Nogueira ED, Olivares FL, Japiassu JC, Vilar C, Vinagre F, Baldani JI, Hemerly AS. 2005. Characterization of glutamine synthetase genes in sugar cane genotypes with different rates of biological nitrogen fixation. Plant. Sci. 169:819–832.

Nylander JAA. 2004. MrModeltest v2. Program distributed by the author. Evolutionary Biology Centre, Uppsala University. https://github.com/nylander/MrModeltest2

Nystedt B, Street NR, Wetterbom A, Zuccolo A, Lin YC, Scofield DG, Vezzi F, Delhomme N, Giacomello S, Alexeyenko A, et al. 2013. The Norway spruce genome sequence and conifer genome evolution. Nature. 497:579–584.

One Thousand Plant Transcriptomes Initiative. 2019. One thousand plant transcriptomes and the phylogenomics of green plants. Nature 574:679–685.

Pascual MB, El-Azaz J, de la Torre FN, Cañas RA, Avila C, Cánovas FM. 2016. Biosynthesis and Metabolic Fate of Phenylalanine in Conifers. Front. Plant Sci. 7:1030.

Pérez-Delgado CM, García-Calderón M, Márquez AJ, Betti M. 2015. Reassimilation of Photorespiratory Ammonium in *Lotus japonicus* Plants Deficient in Plastidic Glutamine Synthetase. PLoS One. 10:e0130438.

Pesole G, Bozzetti MP, Lanave C, Preparata G, Saccone, C. 1991. Glutamine synthetase gene evolution: a good molecular clock. Proc. Nat. Acad. Sci. U.S.A. 88:522–526.

Plett D, Garnett T, Okamoto M. 2017. Molecular genetics to discover and improve nitrogen use efficiency in crop plants. In: Hossain MA, Kamiya T, Burritt DJ, Tran LSP, Fujiwara T, editors. Plant Macronutrient Use Efficiency. London: Academic Press, pp. 93–122.

Raven JA. 2018. Evolution and palaeophysiology of the vascular system and other means of long-distance transport. Philos. Trans. R. Soc. Lond. B Biol. Sci. 373:20160497

Renault H, Werck-Reichhart D, Weng JK. 2019. Harnessing lignin evolution for biotechnological applications. Curr. Opin. Biotechnol. 56:105–111.

Robertson DL, Tartar A. 2006. Evolution of glutamine synthetase in heterokonts: evidence for endosymbiotic gene transfer and the early evolution of photosynthesis. Mol. Biol. Evol. 23:1048–55.

Scott AD, Zimin AV, Puiu D, Workman R, Britton M, Zaman S, Caballero M, Read AC, Bogdanove AJ, Burns E, et al. 2020. A Reference Genome Sequence for Giant Sequoia. G3 (Bethesda) 10:3907–3919.

Shatters RG, Kahn ML. 1989. Glutamine Synthetase II in *Rhizobium:* Reexamination of the Proposed Horizontal Transfer of DNA from Eukaryotes to Prokaryotes. J. Mol. Evol. 29: 422–428.

Sperschneider J, Catanzariti AM, DeBoer K, Petre B, Gardiner DM, Singh KB, Dodds PN, Taylor JM. 2017. LOCALIZER: subcellular localization prediction of both plant and effector proteins in the plant cell. Sci. Rep. 16:44598.

Stevens KA, Wegrzyn JL, Zimin A, Puiu D, Crepeau M, Cardeno C, Paul R, Gonzalez-Ibeas D, Koriabine M, Holtz-Morris AE, et al. 2016. Sequence of the Sugar Pine Megagenome. Genetics, 204:1613–1626.

Tateno Y. 1994. Evolution of glutamine synthetase genes is in accordance with the neutral theory of molecular evolution. Jpn J Genet 69:489–502.

Tegeder M, Masclaux-Daubresse C. 2017. Source and sink mechanisms of nitrogen transport and use. New Phytol. 217:35–53.

Thomsen HC, Erikson D, MØller IS, Schjoerring, JK. 2014. Cytosolic glutamine synthetase: A target for improvement of crop nitrogen use efficiency? Trends Plant Sci. 19:656–663

Wallsgrove RM, Turner JC, Hall NP, Kendall AC, Bright SW. 1987. Barley mutants lacking chloroplast glutamine synthetase-biochemical and genetic analysis. Plant Physiol 83:155–158.

Wan T, Liu ZM, Li LF, Leitch AR, Leitch IL, Lohaus R, Liu ZJ, Xin HP, Huang JL, Li Z, et al. 2018. A genome for gnetophytes and early evolution of seed plants. Nat. Plants 4:82–89.

Ward BB. 2008. Nitrification. In: Jørgensen SE, Fath BD, editors. Encyclopedia of Ecology. Amsterdam (The Netherlands): Elsevier Science. p. 2511–2518.

Wu CS, Chaw SW, Huang YY. 2013. Chloroplast Phylogenomics Indicates that *Ginkgo biloba* Is Sister to Cycads. Genome Biol. Evol. 5:243–254.

Xu G, Fan X, Miller AJ. 2012. Plant Nitrogen Assimilation and Use Efficiency. Annu. Rev. Plant Biol. 63:153–182.

Zimin A, Stevens KA, Crepeau MW, Holtz-Morris A, Koriabine M, Marçais G, Puiu D, Roberts M, Wegrzyn JL, de Jong PJ, et al. 2014. Sequencing and assembly of the 22-gb loblolly pine genome. Genetics, 196:875–90.

