## Supplementary figures and images for "A revised view on the evolution of glutamine synthetase isoenzymes in plants"

### Figure S1

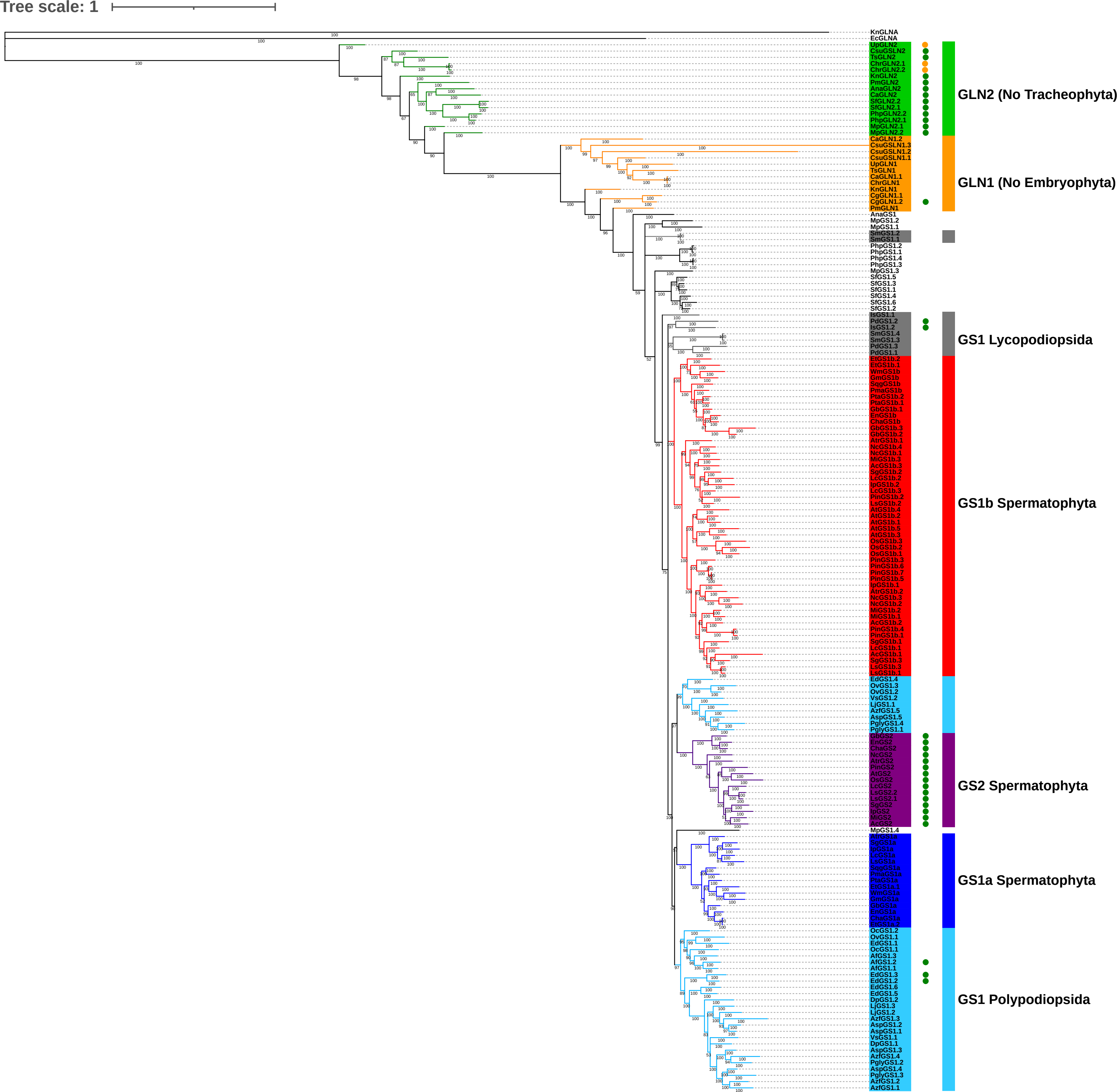

### Figure S2

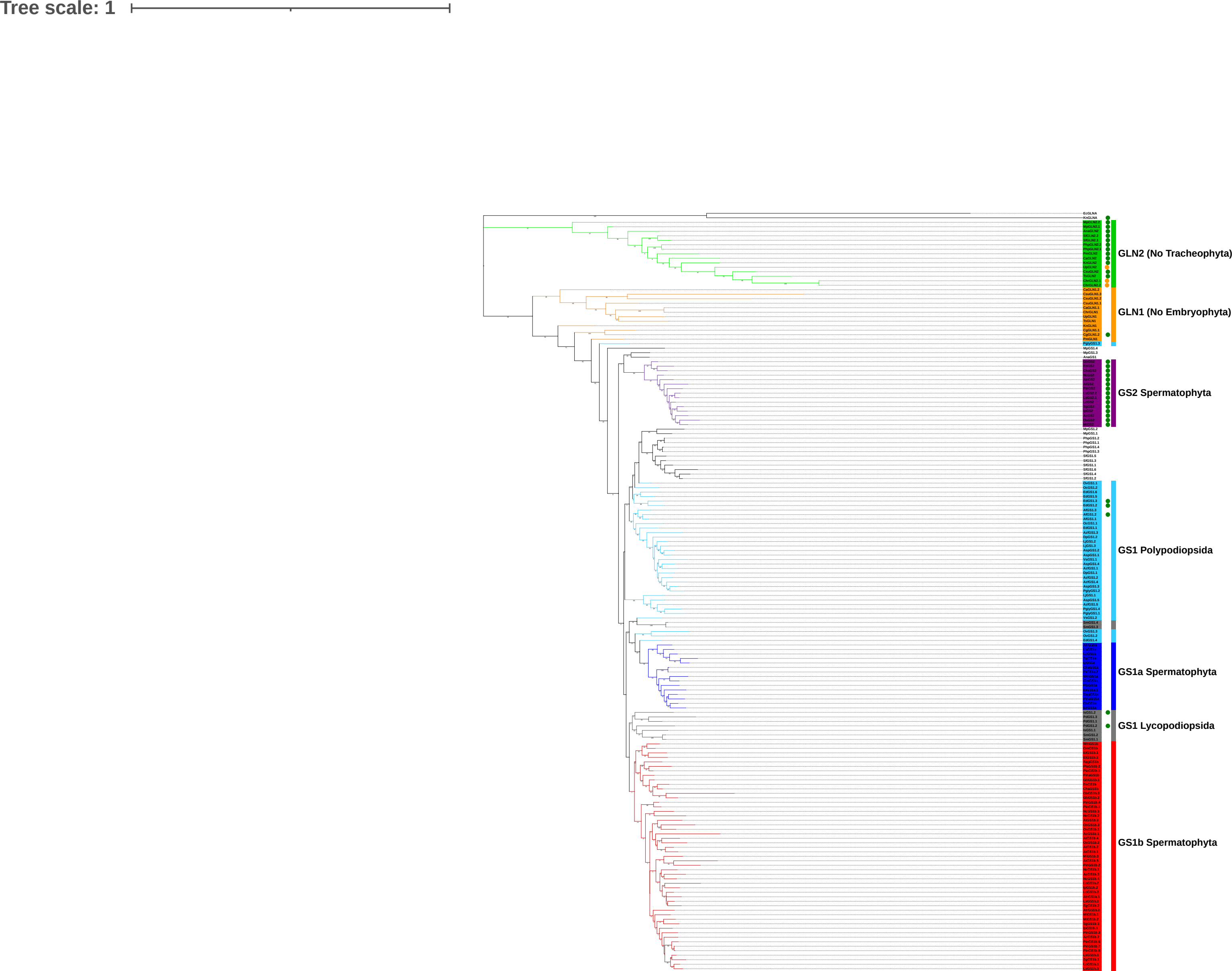
