## Supplementary material for "A revised view on the evolution of glutamine synthetase isoenzymes in plants": Figure S3

```
           1                                                                                                      100
    CrGS2  MSQALVPSLQ WRILPGGVPM TATKMSNCLL PSRGVGLKSA PFNGRLSRSK RSTLHGRNYV SSVRADSVAW NPTENGSASR LIDLLNLDLS PFTDKVIAEY
   ChaGS2  MSQALVPSLQ WRILPGGVPM TATKMSNGLL PSRGVGLKSA PFNGRLSRSK RSTLHGRNYV NSVRADSVAW NPTENGSASR LIDLLNLDLS PFTDKVIAEY
Consensus  MSQALVPSLQ WRILPGGVPM TATKMSNcLL PSRGVGLKSA PFNGRLSRSK RSTLHGRNYV nSVRADSVAW NPTENGSASR LIDLLNLDLS PFTDKVIAEY

           101                                                                                                    200
    CrGS2  LWIGGSGLDI RSKARTVSGP IDNPAKLPKW NYDGSSTGQA PGEDSEVILY PQAIFKDPFR GGNNILVICD SYKPNGEPIP TNKRANAAKI FSQKKVIDEE
   ChaGS2  LWIGGSGLDI RSKARTVSGP IDNPAKLPKW NYDGSSTGQA PGEDSEVILY PQAIFKDPFR GGNNILVICD CYKPNGEPIP TNKRANAAKI FSQKKVIDEE
Consensus  LWIGGSGLDI RSKARTVSGP IDNPAKLPKW NYDGSSTGQA PGEDSEVILY PQAIFKDPFR GGNNILVICD cYKPNGEPIP TNKRANAAKI FSQKKVIDEE

           201                                                                                                    300
    CrGS2  PWYGIEQEYT LLQKNVKWPL GWPIGGYPGP QGPYYCGTGV DKAYGRVIAD AHYKACVYAG IKVSGINSEV MPGQWEYQVG PSVGIASGDH LWCSRYILER
   ChaGS2  PWYGIEQEYT LLQKNVKWPL GWPIGGYPGP QGPYYCGTGV DKAYGRVIAD AHYKACVYAG IKVSGINSEV MPGQWEYQVG PSVGIASGDH LWCSRYILER
Consensus  PWYGIEQEYT LLQKNVKWPL GWPIGGYPGP QGPYYCGTGV DKAYGRVIAD AHYKACVYAG IKVSGINSEV MPGQWEYQVG PSVGIASGDH LWCSRYILER

           301                                                                                                    400
    CrGS2  ITEMAGVVLS LDPKPIEGDW NGAGCHTNYS TKSMREDGGY EVIKKAILNL GLRHKEHISA YGEGNERRLT GHHETANINA FSWGVANRGA SIRVGRETEK
   ChaGS2  ITEMAGVVLS LDPKPIDGDW NGAGCHTNYS TKSMREDGGY EVIKKAILNL GLRHKEHISA YGEGNERRLT GHHETANINA FSWGVANRGA SIRVGRETEK
Consensus  ITEMAGVVLS LDPKPI#GDW NGAGCHTNYS TKSMREDGGY EVIKKAILNL GLRHKEHISA YGEGNERRLT GHHETANINA FSWGVANRGA SIRVGRETEK

           401                                          449
    CrGS2  QGKGYLEDRR PASNMDPYVV TSMLAETTIL WEPAPEAGTH AAKELQLQI
   ChaGS2  QGKGYLEDRR PASNMDPYVV TSMLAETTIL WEPAPEAGTH AAKELQLQI
Consensus  QGKGYLEDRR PASNMDPYVV TSMLAETTIL WEPAPEAGTH AAKELQLQI
```
