## Supplementary material for "A revised view on the evolution of glutamine synthetase isoenzymes in plants": Figure S4

```
           1                                                                                                       100
     CrGS2 ATGTCGCAGG CATTGGTACC GTCGTTGCAA TGGCGAATTT TGCCGGGAGG TGTCCCCATG ACGGCAACGA AGATGTCAAA TTGTTTGCTT CCTAGCCGGG
    ChaGS2 ATGTCGCAGG CATTGGTACC GTCGTTGCAA TGGCGAATTT TGCCGGGAGG TGTCCCTATG ACGGCAACGA AGATGTCAAA TGGTTTGCTT CCTAGCCGGG
 Consensus ATGTCGCAGG CATTGGTACC GTCGTTGCAA TGGCGAATTT TGCCGGGAGG TGTCCCcATG ACGGCAACGA AGATGTCAAA TgGTTTGCTT CCTAGCCGGG

           101                                                                                                     200
     CrGS2 GAGTTGGTTT GAAGAGTGCA CCATTCAACG GGAGACTTAG TAGATCGAAG CGTAGCACTC TGCATGGTAG GAATTATGTC AGCAGTGTTC GCGCCGATTC
    ChaGS2 GAGTTGGTTT GAAGAGTGCA CCATTCAACG GGAGACTTAG TAGATCGAAG CGTAGCACTC TGCATGGTAG GAATTATGTC AACAGTGTTC GCGCCGATTC
 Consensus GAGTTGGTTT GAAGAGTGCA CCATTCAACG GGAGACTTAG TAGATCGAAG CGTAGCACTC TGCATGGTAG GAATTATGTC AaCAGTGTTC GCGCCGATTC

           201                                                                                                     300
     CrGS2 CGTTGCTTGG AATCCGACTG AAAACGGAAG CGCATCACGA CTGATTGATC TCCTCAACTT AGATTTATCT CCATTTACTG ACAAAGTCAT TGCAGAGTAT
    ChaGS2 CGTTGCTTGG AATCCGACTG AAAACGGAAG CGCATCACGA CTGATTGATC TCCTCAACTT AGATTTATCT CCATTTACTG ACAAAGTCAT TGCAGAATAT
 Consensus CGTTGCTTGG AATCCGACTG AAAACGGAAG CGCATCACGA CTGATTGATC TCCTCAACTT AGATTTATCT CCATTTACTG ACAAAGTCAT TGCAGAaTAT

           301                                                                                                     400
     CrGS2 TTATGGATTG GAGGATCGGG ATTGGACATT CGTAGCAAGG CTAGAACAGT AAGTGGTCCC ATCGACAACC CTGCCAAGCT TCCAAAATGG AACTACGATG
    ChaGS2 TTATGGATTG GAGGATCGGG ATTGGATATT CGTAGCAAGG CTAGAACAGT AAGTGGTCCC ATCGACAACC CTGCCAAGCT TCCAAAATGG AACTACGATG
 Consensus TTATGGATTG GAGGATCGGG ATTGGAcATT CGTAGCAAGG CTAGAACAGT AAGTGGTCCC ATCGACAACC CTGCCAAGCT TCCAAAATGG AACTACGATG

           401                                                                                                     500
     CrGS2 GATCCAGCAC GGGTCAAGCT CCAGGGGAGG ACAGTGAAGT CATTCTCTAT CCACAGGCCA TCTTCAAGGA CCCTTTCCGA GGAGGAAACA ACATATTGGT
    ChaGS2 GATCCAGCAC GGGTCAAGCT CCAGGGGAGG ACAGTGAAGT CATTCTCTAT CCACAGGCGA TCTTCAAGGA CCCTTTCCGA GGAGGAAACA ACATATTGGT
 Consensus GATCCAGCAC GGGTCAAGCT CCAGGGGAGG ACAGTGAAGT CATTCTCTAT CCACAGGCcA TCTTCAAGGA CCCTTTCCGA GGAGGAAACA ACATATTGGT

           501                                                                                                     600
     CrGS2 AATATGCGAT TCTTACAAGC CAAATGGGGA GCCAATTCCT ACAAACAAGC GTGCCAATGC CGCTAAGATT TTCAGCCAGA AAAAAGTCAT TGATGAGGAG
    ChaGS2 AATATGCGAT TGTTACAAGC CAAATGGGGA GCCAATTCCT ACAAACAAGC GTGCCAATGC CGCTAAGATT TTCAGCCAGA AAAAAGTCAT TGATGAGGAG
 Consensus AATATGCGAT TcTTACAAGC CAAATGGGGA GCCAATTCCT ACAAACAAGC GTGCCAATGC CGCTAAGATT TTCAGCCAGA AAAAAGTCAT TGATGAGGAG

           601                                                                                                     700
     CrGS2 CCTTGGTATG GCATCGAGCA AGAGTACACT TTGTTGCAGA AGAATGTTAA ATGGCCTCTT GGATGGCCCA TTGGAGGTTA CCCGGGACCA CAGGGTCCGT
    ChaGS2 CCTTGGTATG GCATCGAGCA AGAGTACACT TTGTTGCAGA AGAATGTTAA ATGGCCTCTT GGATGGCCCA TTGGAGGTTA CCCGGGACCA CAGGGTCCGT
 Consensus CCTTGGTATG GCATCGAGCA AGAGTACACT TTGTTGCAGA AGAATGTTAA ATGGCCTCTT GGATGGCCCA TTGGAGGTTA CCCGGGACCA CAGGGTCCGT

           701                                                                                                     800
     CrGS2 ATTATTGTGG AACGGGAGTA GACAAAGCCT ACGGCCGTGT CATCGCAGAC GCTCATTATA AGGCTTGTGT TTATGCTGGA ATCAAAGTTA GCGGCATCAA
    ChaGS2 ATTATTGTGG AACGGGAGTA GACAAAGCCT ACGGCCGTGT CATCGCAGAC GCTCATTATA AGGCTTGTGT TTATGCTGGA ATCAAAGTTA GCGGCATCAA
 Consensus ATTATTGTGG AACGGGAGTA GACAAAGCCT ACGGCCGTGT CATCGCAGAC GCTCATTATA AGGCTTGTGT TTATGCTGGA ATCAAAGTTA GCGGCATCAA

           801                                                                                                     900
     CrGS2 TAGCGAGGTT ATGCCCGGAC AGTGGGAGTA CCAAGTGGGT CCCAGCGTGG GAATTGCTTC CGGAGATCAT CTCTGGTGTT CTCGTTACAT TCTTGAGAGA
    ChaGS2 TAGCGAGGTT ATGCCCGGAC AGTGGGAGTA CCAAGTGGGT CCCAGCGTGG GAATTGCTTC CGGAGATCAT CTCTGGTGTT CTCGTTACAT TCTTGAGAGA
 Consensus TAGCGAGGTT ATGCCCGGAC AGTGGGAGTA CCAAGTGGGT CCCAGCGTGG GAATTGCTTC CGGAGATCAT CTCTGGTGTT CTCGTTACAT TCTTGAGAGA

           901                                                                                                    1000
     CrGS2 ATTACAGAAA TGGCAGGCGT GGTGCTCTCG CTCGATCCGA AGCCCATAGA GGGTGATTGG AATGGTGCGG GCTGTCACAC TAATTATAGC ACTAAGTCCA
    ChaGS2 ATTACAGAAA TGGCAGGCGT GGTGCTCTCG CTCGATCCGA AGCCCATAGA TGGTGATTGG AATGGTGCGG GCTGTCACAC TAATTATAGC ACTAAGTCCA
 Consensus ATTACAGAAA TGGCAGGCGT GGTGCTCTCG CTCGATCCGA AGCCCATAGA gGGTGATTGG AATGGTGCGG GCTGTCACAC TAATTATAGC ACTAAGTCCA

           1001                                                                                                   1100
     CrGS2 TGCGAGAGGA TGGGGGTTAT GAGGTGATAA AGAAAGCTAT ATTAAATCTT GGGTTGCGCC ATAAGGAACA CATCAGTGCC TATGGAGAAG GAAATGAAAG
    ChaGS2 TGCGAGAGGA TGGGGGTTAT GAGGTGATAA AGAAAGCTAT ATTAAATCTT GGGTTGCGCC ATAAGGAACA CATCAGTGCC TATGGAGAAG GAAATGAAAG
 Consensus TGCGAGAGGA TGGGGGTTAT GAGGTGATAA AGAAAGCTAT ATTAAATCTT GGGTTGCGCC ATAAGGAACA CATCAGTGCC TATGGAGAAG GAAATGAAAG

           1101                                                                                                   1200
     CrGS2 AAGATTAACT GGACACCATG AGACTGCAAA CATCAATGCA TTCTCTTGGG GTGTAGCAAA TCGTGGTGCA TCCATTCGCG TTGGACGTGA AACAGAGAAG
    ChaGS2 AAGATTAACT GGACACCATG AGACTGCAAA CATCAATGCA TTCTCTTGGG GTGTAGCAAA TCGTGGTGCA TCCATTCGCG TTGGACGTGA AACAGAGAAG
 Consensus AAGATTAACT GGACACCATG AGACTGCAAA CATCAATGCA TTCTCTTGGG GTGTAGCAAA TCGTGGTGCA TCCATTCGCG TTGGACGTGA AACAGAGAAG

           1201                                                                                                   1300
     CrGS2 CAAGGAAAAG GCTACCTGGA GGATAGGCGG CCTGCTTCGA ACATGGATCC ATATGTCGTG ACTTCCATGC TGGCAGAAAC TACAATCCTA TGGGAACCGG
    ChaGS2 CAAGGAAAAG GCTACCTGGA GGATAGGCGG CCTGCTTCGA ACATGGATCC ATATGTCGTG ACTTCCATGC TGGCAGAAAC TACAATCCTG TGGGAACCGG
 Consensus CAAGGAAAAG GCTACCTGGA GGATAGGCGG CCTGCTTCGA ACATGGATCC ATATGTCGTG ACTTCCATGC TGGCAGAAAC TACAATCCTa TGGGAACCGG

           1301                                              1350
     CrGS2 CTCCCGAGGC AGGAACTCAC GCTGCCAAGG AGTTACAGTT GCAAATTTGA
    ChaGS2 CTCCCGAGGC AGGAACTCAC GCTGCCAAGG AGTTACAGTT GCAAATTTGA
 Consensus CTCCCGAGGC AGGAACTCAC GCTGCCAAGG AGTTACAGTT GCAAATTTGA
```
