## Supplementary material for "A revised view on the evolution of glutamine synthetase isoenzymes in plants": Figure S5

```
                      ..500   ▼    511..  ..583   ▼    594..
  1 EcGLNA          ...NGSGMHCHMSLS...  ...SARNRSASIRIP...
  2 ChrGLN1         ...NGSG--GHTNYS...  ...GVANRGCSIRVG...
  3 ChrGLN2.2       ...NGTG--AHTNFS...  ...GVADRGSSIRIP...
  4 ChrGLN2.1       ...NGTG--AHTNFS...  ...GVADRGSSIRIP...
  5 CsuGLN1.1       ...NGAG--GHTNFS...  ...GVANRGASIRVG...
  6 CsuGLN2         ...NGTG--AHTNFS...  ...GVADRGSSIRIP...
  7 CsuGLN1.2       ...AGTG--GHTNYS...  ...GVGDRGASVRVG...
  8 CsuGLN1.3       ...SGNG--AAVKFS...  ...GMENRNASIRIP...
  9 TsGLN2          ...NGTG--AHTNYS...  ...GFADRGASIRIP...
 10 TsGLN1          ...NGAG--GHTNYS...  ...GVANRGCSIRVG...
 11 UpGLN1          ...NGAG--GHTNYS...  ...GVANRGCSIRIG...
 12 UpGLN2          ...NGAG--AHTNYS...  ...GVADRGSSIRIP...
 13 CaGLN1.2        ...NGAG--CHTNYS...  ...GVADRGKSIRVG...
 14 CaGLN2          ...NGTG--GHTNYS...  ...GVADRGASIRIP...
 15 CaGLN1.1        ...NGSG--GHTNYS...  ...GVANRGCSIRVG...
 16 KnGLN2          ...NGTG--AHTNYS...  ...GVSDRGASIRIP...
 17 KnGLN1          ...NGAG--CHTNYS...  ...GVANRGASIRVG...
 18 KnGLNA          ...AGSS--CHVHMS...  ...SKDNRTAPFRI-...
 19 CgGLN1.2        ...NGAG--CHTNYS...  ...GVANRGASIRVG...
 20 CgGLN1.1        ...NGAG--CHTNYS...  ...GVANRGASIRVG...
 21 PmGLN1          ...NGAG--AHTNYS...  ...GVANRGCSIRVG...
 22 PmGLN2          ...NGTG--AHTNYS...  ...GVADRGASIRIP...
 23 AnaGS1          ...NGAG--CHTNYS...  ...GVANRGASIRVG...
 24 AnaGLN2         ...NGAG--AHTNYS...  ...GVADRGASIRIP...
 25 MpGS1.1         ...NGAG--CHTNFS...  ...GVANRGSSIRVG...
 26 MpGS1.2         ...NGAG--CHTNFS...  ...GVANRGASIRVG...
 27 MpGLN2.1        ...NGAG--AHTNYS...  ...GVADRGASIRIP...
 28 MpGLN2.2        ...NGAG--AHTNFS...  ...GVADRGASVRIP...
 29 MpGS1.3         ...NGAG--CHTNYS...  ...GVANRGASVRVG...
 30 MpGS1.4         ...NGAG--CHTNYS...  ...GVADRGASIRVG...
 31 SfGLN2.1        ...NGAG--AHTNYS...  ...GVADRGASIRIP...
 32 SfGLN2.2        ...NGAG--AHTNYS...  ...GVADRGASIRIP...
 33 SfGS1.1         ...NGAG--CHTNYS...  ...GVANRGASVRVG...
 34 SfGS1.2         ...NGAG--CHTNYS...  ...GVANRGASIRVG...
 35 SfGS1.3         ...NGAG--CHTNYS...  ...GVANRGASIRVG...
 36 SfGS1.4         ...NGAG--CHTNYS...  ...GVANRGASIRVG...
 37 SfGS1.5         ...NGAG--CHTNYS...  ...GVANRGASIRVG...
 38 SfGS1.6         ...NGAG--CHANYS...  ...GVGNRGVSIRVG...
 39 PhpGS1.1        ...NGAG--CHTNYS...  ...GVANRGASVRVG...
 40 PhpGS1.2        ...NGAG--CHTNYS...  ...GVANRGASVRVG...
 41 PhpGS1.3        ...NGAG--CHTNYS...  ...GVANRGASVRVG...
 42 PhpGS1.4        ...NGAG--CHTNYS...  ...GVANRGASVRVG...
 43 PhpGLN2.1       ...NGAG--AHTNYS...  ...GVADRGASIRIP...
 44 PhpGLN2.2       ...NGAG--AHTNYS...  ...GVADRGASIRIP...
 45 SmGS1.1         ...NGAG--CHANYS...  ...GVANRGASVRVG...
 46 SmGS1.2         ...NGAG--CHTNYS...  ...GVANRGASVRVG...
 47 SmGS1.3         ...NGAG--AHTNYS...  ...GVANRGASVRVG...
 48 SmGS1.4         ...NGAG--AHTNYS...  ...GVANRGASVRVG...
 49 IsGS1.2         ...NGAG--AHTNYS...  ...GVANRGASIRVG...
 50 IsGS1.1         ...NGAG--AHTNYS...  ...GVANRGASVRVG...
 51 PdGS1.1         ...NGAG--AHTNYS...  ...GVANRGASIRVG...
 52 PdGS1.2         ...NGAG--AHTNYS...  ...GVANRGASIRVG...
 53 PdGS1.3         ...NGAG--AHVNYS...  ...GAGTRSTSVRVS...
 54 EdGS1.1         ...NGAG--CHTNYS...  ...GVANRGASIRVG...
 55 EdGS1.5         ...NGAG--CHTNYS...  ...GVANRGASVRVG...
 56 EdGS1.6         ...NGAG--CHTNYS...  ...GVANRGASIRVG...
 57 EdGS1.2         ...NGAG--CHTNYS...  ...GVANRGSSIRVG...
 58 EdGS1.3         ...NGAG--CHTNFS...  ...GVANRGSSIRVG...
 59 EdGS1.4         ...NGAG--CHTNYS...  ...GVANRGASVRVG...
 60 AfGS1.1         ...NGAG--CHTNYS...  ...GVANRGASIRVG...
 61 AfGS1.3         ...NGAG--CHTNYS...  ...GVANRGASIRVG...
 62 AfGS1.2         ...NGAG--CHTNYS...  ...GVANRGASVRVG...
 63 OvGS1.1         ...NGAG--CHTNYS...  ...GVANRGASIRVG...
 64 OvGS1.2         ...NGAG--CHANYS...  ...GVANRGASVRVG...
 65 OvGS1.3         ...NGGG--CHTNYS...  ...GVANRGTSIRIG...
 66 PglyGS1.1       ...NGAG--CHTNFS...  ...GVADRGASIRVG...
 67 PglyGS1.2       ...NGAG--CHTNYS...  ...GVANRGASIRVG...
 68 PglyGS1.3       ...NGAG--CHTNYS...  ...GVANRGASIRVG...
 69 PglyGS1.4       ...NGAG--CHTNFS...  ...GVANRGASVRVG...
 70 DpGS1.1         ...NGAG--CHTNYS...  ...GVANRGASIRVG...
 71 DpGS1.2         ...NGAG--CHTNYS...  ...GVANRGASIRVG...
 72 VsGS1.1         ...NGAG--CHTNYS...  ...GVANRGASVRVG...
 73 VsGS1.2         ...NGAG--CHTNYS...  ...GVANRGASVRVG...
 74 OcGS1.1         ...NGAG--CHTNYS...  ...GVANRGASIRVG...
 75 OcGS1.2         ...NGAG--CHTNYS...  ...GVANRGASIRVG...
 76 AzfGS1.1        ...NGAG--CHTNYS...  ...GVANRGASIRVG...
 77 AzfGS1.2        ...NGAG--CHTNYS...  ...GVANRGASIRVG...
 78 AzfGS1.3        ...NGAG--CHTNYS...  ...GVANRGVSVRVG...
 79 AzfGS1.4        ...NGAG--CHTNYS...  ...GVANRGASIRVG...
 80 AzfGS1.5        ...NGAG--CHTNFS...  ...GVANRGASVRVG...
 81 LjGS1.1         ...NGAG--CHTNYS...  ...GVANRGASVRVG...
 82 LjGS1.2         ...NGAG--CHTNYS...  ...GVANRGASIRVG...
 83 LjGS1.3         ...NGAG--CHTNYS...  ...GVANRGASIRVG...
 84 AspGS1.1        ...NGAG--CHTNYS...  ...GVANRGASIRVG...
 85 AspGS1.2        ...NGAG--CHTNYS...  ...GVANRGASIRVG...
 86 AspGS1.3        ...NGAG--CHTNYS...  ...GVANRGASIRVG...
 87 AspGS1.4        ...NGAG--CHTNYS...  ...GVADRGASIRVG...
 88 AspGS1.5        ...NGAG--CHTNYS...  ...GVANRGASVRVG...
 89 GmGS1b          ...NGAG--AHTNYS...  ...GVANRGASVRVG...
 90 GmGS1a          ...NGAG--CHSNYS...  ...GVANRGASVRVG...
 91 WmGS1b          ...NGAG--AHTNYS...  ...GVANRGASVRVG...
 92 WmGS1a          ...NGAG--CHANYS...  ...GVANRGASVRVG...
 93 EtGS1b.2        ...NGAG--AHTNYS...  ...GVANRGASVRVG...
 94 EtGS1b.1        ...NGAG--AHTNYS...  ...GVANRGASVRVG...
 95 EtGS1a.1        ...NGAG--CHTNYS...  ...GVANRGASVRVG...
 96 EtGS1a.2        ...NGAG--CHTNYS...  ...GVANRGASVRVG...
 97 PmaGS1b         ...NGAG--AHTNYS...  ...GVANRGASIRVG...
 98 PmaGS1a         ...NGAG--CHTNYS...  ...GVANRGASVRVG...
 99 SqgGS1b         ...NGAG--AHTNYS...  ...GVANRGASIRVG...
100 SqgGS1a         ...NGAG--CHTNYS...  ...GVANRGASIRVG...
101 PtaGS1a         ...NGAG--CHTNYS...  ...GVANRGASVRVG...
102 PtaGS1b.1       ...NGAG--AHTNYS...  ...GVANRGASIRVG...
103 PtaGS1b.2       ...NGAG--AHTNYS...  ...GVANRGASVRIG...
104 ChaGS1a         ...NGAG--CHTNYS...  ...GVANRGASVRVG...
105 ChaGS1b         ...NGAG--AHANYS...  ...GVANRGASIRVG...
106 ChaGS2          ...NGAG--CHTNYS...  ...GVANRGASIRVG...
107 EnGS1a          ...NGAG--CHTNYS...  ...GVANRGASIRVG...
108 EnGS2           ...NGAG--CHTNYS...  ...GVANRGASIRVG...
109 EnGS1b          ...NGAG--AHANYS...  ...GVANRGASVRVG...
110 GbGS1a          ...NGAG--CHTNYS...  ...GVANRGASVRVG...
111 GbGS1b.1        ...NGAG--AHTNYS...  ...GVANRGASIRVG...
112 GbGS1b.2        ...NGAG--AHTNYN...  ...GVADRGASIRVG...
113 GbGS1b.3        ...KGGR--AHTNYS...  ...GVAKREASIRAG...
114 GbGS2           ...NGAG--CHTNYS...  ...GVANRGASIRVG...
115 AtrGS1a         ...NGAG--CHTNYS...  ...GVANRGASVRVG...
116 AtrGS1b.1       ...NGAG--AHTNYS...  ...GVANRGASVRVG...
117 AtrGS1b.2       ...NGAG--CHSNYS...  ...GVANRGASIRVG...
118 AtrGS2          ...NGAG--CHTNFS...  ...GVANRGCSIRVG...
119 NcGS1b.1        ...NGAG--AHTNYS...  ...GVANRGASVRIG...
120 NcGS1b.2        ...NGAG--AHTNYS...  ...GVANRGASIRVG...
121 NcGS1b.3        ...NGAG--AHTNYS...  ...GVANRGASIRVG...
122 NcGS1b.4        ...NGAG--AHTNYS...  ...GVANRGASIRVG...
123 NcGS2           ...NGAG--CHTNYS...  ...GVANRGCSIRVG...
124 IpGS1b.2        ...NGAG--AHTNYS...  ...GVANRGASIRVG...
125 IpGS2           ...NGAG--CHTNYS...  ...GVANRGCSIRVG...
126 IpGS1b.1        ...NGAG--AHTNYS...  ...GVANRGASIRVG...
127 IpGS1a          ...NGAG--CHTNYS...  ...GVANRGASIRVG...
128 SgGS1b.1        ...NGAG--AHTNYS...  ...GVANRGASIRVG...
129 SgGS1b.2        ...NGAG--AHTNYS...  ...GVANRGASIRVG...
130 SgGS1a          ...NGAG--CHTNYS...  ...GVANRGASIRVG...
131 SgGS2           ...NGAG--CHTNYS...  ...GVANRGCSIRVG...
132 SgGS1b.3        ...NGAG--AHTNYS...  ...GVANRGASIRVG...
133 LsGS1a          ...NGAG--CHTNYS...  ...GVANRGASIRVG...
134 LsGS1b.1        ...NGAG--AHTNYS...  ...GVANRGASIRVG...
135 LsGS1b.2        ...NGAG--AHTNYS...  ...GVANRGASIRVG...
136 LsGS1b.3        ...NGAG--AHTNYS...  ...GVANRGASIRVG...
137 LsGS2.1         ...NGAG--CHTNYS...  ...GVANRGCSIRVG...
138 LsGS2.2         ...NGAG--CHTNYS...  ...GVANRGCSIRVG...
139 PinGS1b.1       ...NGAG--AHTNYS...  ...GVANRGASIRVG...
140 PinGS1b.2       ...NGAG--AHTNYS...  ...GVANRGASVRIG...
141 PinGS1b.3       ...NGAG--AHTNYS...  ...GVANRGASIRVG...
142 PinGS2          ...NGAG--CHTNYS...  ...GVANRGCSIRVG...
143 PinGS1b.4       ...NGAG--AHTNYS...  ...GVANRGASIRVG...
144 PinGS1b.5       ...NGAG--AHTNYS...  ...GVANRGASIRVG...
145 PinGS1b.6       ...NGAG--AHTNYS...  ...GVANRGASVRVS...
146 PinGS1b.7       ...NGAG--AHTNYS...  ...GVANRGASIRVG...
147 LcGS1a          ...NGAG--CHTNYS...  ...GVANRGASIRVG...
148 LcGS1b.1        ...NGAG--AHTNYS...  ...GVANRGASIRVG...
149 LcGS1b.2        ...NGAG--AHTNYS...  ...GIRNRAASIRVG...
150 LcGS1b.3        ...NGAG--AHTNYS...  ...GVANRGASIRVG...
151 LcGS2           ...NGAG--CHTNYS...  ...GVANRGCSIRVG...
152 AcGS1b.1        ...NGTG--AHTNFS...  ...GVGNRAASIRVG...
153 AcGS1b.2        ...NGAG--AHCNYS...  ...GVANRGASVRVG...
154 AcGS1b.3        ...NGAG--AHTNYS...  ...GVANRGASIRVG...
155 AcGS2           ...NGAG--CHTNYS...  ...GVANRGCSIRVG...
156 MiGS1b.1        ...NGAG--AHTNYS...  ...GVADRGASIRVG...
157 MiGS1b.2        ...NGAG--AHTNYS...  ...GVANRGASIRVG...
158 MiGS1b.3        ...NGAG--AHTNYS...  ...GVANRGASIRVG...
159 MiGS2           ...NGAG--CHTNYS...  ...GVANRGCSIRVG...
160 OsGS1b.1        ...NGAG--AHTNYS...  ...GVANRGASVRVG...
161 OsGS1b.2        ...NGAG--AHTNFS...  ...GVANRGASIRVG...
162 OsGS1b.3        ...NGAG--AHTNYS...  ...GVANRGASVRVG...
163 OsGS2           ...NGAG--CHTNYS...  ...GVANRGCSIRVG...
164 AtGS1b.1        ...NGAG--AHCNYS...  ...GVANRGASIRVG...
165 AtGS1b.2        ...NGAG--AHTNYS...  ...GVANRGASIRVG...
166 AtGS1b.3        ...NGAG--AHCNYS...  ...GVANRGASVRVG...
167 AtGS1b.4        ...NGAG--AHTNYS...  ...GVANRGASIRVG...
168 AtGS1b.5        ...NGAA--AHTNFS...  ...GVADRGASVRVG...
169 AtGS2           ...NGAG--CHTNYS...  ...GVANRGCSIRVG...
```
