## Supplementary material for "A revised view on the evolution of glutamine synthetase isoenzymes in plants": Table S2

**Table S2.** List of primers used for RT-qPCR experiments *CrGS2* cloning.

| **Gene** | **Organism** | **Primer name** | **Sequence (5’ → 3’)** | **Primer sense** |
| --- | --- | --- | --- | --- |
| ***GbGS1b.1*** | *Ginkgo biloba* | Gb_GS1b.1-F | GATTGTTTGCCATGAATGAGGGT | Forward |
| ***GbGS1b.1*** | *Ginkgo biloba* | Gb_GS1b.1-R | ATTCCACAAACAATGCAGCATC | Reverse |
| ***GbGS1b.2*** | *Ginkgo biloba* | Gb_GS1b.2-F | TCTCGTTGAATAAGCAGACCACA | Forward |
| ***GbGS1b.2*** | *Ginkgo biloba* | Gb_GS1b.2-R | ATTGTTCTCTTCTCAAACGCGC | Reverse |
| ***GbGS1b.3*** | *Ginkgo biloba* | Gb_GS1b.3-F | GCTTGTGGAATAATGAGGTTCGT | Forward |
| ***GbGS1b.3*** | *Ginkgo biloba* | Gb_GS1b.3-R | GTCTCCGCAAACAGGCAATATC | Reverse |
| ***GbGS1a*** | *Ginkgo biloba* | Gb_GS1a-F | GTCGCCCTGCATCCAACATG | Forward |
| ***GbGS1a*** | *Ginkgo biloba* | Gb_GS1a-R | TGAAGCACTCACGATTATGGCT | Reverse |
| ***GbGS2*** | *Ginkgo biloba* | Gb_GS2-F | ACTAACTGGAAAGCACGAGACT | Forward |
| ***GbGS2*** | *Ginkgo biloba* | Gb_GS2-R | GCCTTTTCCTTGCTTCTCTGTT | Reverse |
| ***Saposin-like aspartyl protease*** | *Ginkgo biloba* | Gb_1135-F | AGTGTTCTTGTTCGACATTCCA | Forward |
| ***Saposin-like aspartyl protease*** | *Ginkgo biloba* | Gb_1135-R | GCATTTGTAGCCAGCACTTTGA | Reverse |
| ***Myosin heavy chain-related*** | *Ginkgo biloba* | Gb_13291-F | TGAAAGCAGAAGTCTCCACATGA | Forward |
| ***Myosin heavy chain-related*** | *Ginkgo biloba* | Gb_13291-R | CAATGCTTCCCTTGCCTCAAAC | Reverse |
| ***RNA binding protein*** | *Ginkgo biloba* | Gb_27526-F | GAACACAAGCAAATCTCCTCTTTC | Forward |
| ***RNA binding protein*** | *Ginkgo biloba* | Gb_27526-R | TGCATCTATCGTGACCCAAGAG | Reverse |
| ***MgGS1b.1*** | *Magnolia grandiflora* | Mg_GS1b.1-F | GCCCACCACATTATCTTCTTCA | Forward |
| ***MgGS1b.1*** | *Magnolia grandiflora* | Mg_GS1b.1-R | TTGCCTCACAGACCCACTAAAA | Reverse |
| ***MgGS1b.2*** | *Magnolia grandiflora* | Mg_GS1b.2-F | TGAAGGACTTACAAAATACACCGG | Forward |
| ***MgGS1b.2*** | *Magnolia grandiflora* | Mg_GS1b.2-R | GCAGCCACCCAGTTTGAAAATA | Reverse |
| ***MgGS1b.3*** | *Magnolia grandiflora* | Mg_GS1b.3-F | GGGCCTGAATTTCCATTGGTTT | Forward |
| ***MgGS1b.3*** | *Magnolia grandiflora* | Mg_GS1b.3-R | CCTGCCTCTCTACAACCTCAAA | Reverse |
| ***MgGS1a*** | *Magnolia grandiflora* | Mg_GS1a-F | TGTCAAGAGAAAGCTTCCCTGT | Forward |
| ***MgGS1a*** | *Magnolia grandiflora* | Mg_GS1a-R | CAACACTCAAGGTAAATGCCCA | Reverse |
| ***MgGS2*** | *Magnolia grandiflora* | Mg_GS2-F | TGACCCACCCCTTCTATATTATCA | Forward |
| ***MgGS2*** | *Magnolia grandiflora* | Mg_GS2-R | TGCCAATCAAACAAACACCCA | Reverse |
| ***Actin*** | *Magnolia grandiflora* | Mg_Actin-F | CCATCACCGGAATCAAGCACAATA | Forward |
| ***Actin*** | *Magnolia grandiflora* | Mg_Actin-R | CAAGGCCAACAGGGAGAAAATGAC | Reverse |
| ***RNA binding protein*** | *Magnolia grandiflora* | Mg_27526-F | TGATGGCTGATAATTGGTGGTGA | Forward |
| ***RNA binding protein*** | *Magnolia grandiflora* | Mg_27526-R | GGCTATGTGATGGAGTGGGTC | Reverse |
| ***PpGS1a*** | *Pinus pinaster* | Pp_GS1a-F | ATCGAGGAGCTTCAGTTAGAG | Forward |
| ***PpGS1a*** | *Pinus pinaster* | Pp_GS1a-R | TGGTCGTCTCAGCAATCATAGA | Reverse |
| ***PpGS1b*** | *Pinus pinaster* | Pp_GS1b-F | CCCAATTGTTTGTGGGGGATA | Forward |
| ***PpGS1b*** | *Pinus pinaster* | Pp_GS1b-F | CTGAATGACAAACTAGACACTG | Reverse |
| ***Saposin-like aspartyl protease*** | *Pinus pinaster* | Pp_1135-F | AGTATGCTAAGGAATCGTGCCT | Forward |
| ***Saposin-like aspartyl protease*** | *Pinus pinaster* | Pp_1135-F | GTCCATAATTACACACGAACAGA | Reverse |
| ***CrGS2*** | *Cyscas revoluta* | GS2_F4 | TCGGGATTGGACATTCGTAGCAAGGCTAGA | Forward |
| ***CrGS2*** | *Cyscas revoluta* | GS2_R5 | AGCGAGAGCACCACGCCTGCCAT | Reverse |
| ***CrGS2*** | *Cyscas revoluta* | GS2_F9 | ATGTCGCAGGCATTGGTA | Forward |
| ***CrGS2*** | *Cyscas revoluta* | GS2_R4 | TCTAGCCTTGCTACGAATGTCCAATCCCGA | Reverse |
